# Effects of diffusion MRI spatial resolution on human brain short-range association fiber reconstruction and structural connectivity estimation

**DOI:** 10.1101/2025.06.04.657810

**Authors:** Jialan Zheng, Ziyu Li, Wen Zhong, Ziang Wang, Zihan Li, Hongjia Yang, Mingxuan Liu, Xiaozhi Cao, Congyu Liao, David H. Salat, Susie Y. Huang, Qiyuan Tian

## Abstract

Short-range association fibers (SAFs) are critical for cortical communications but are often underestimated in conventional resolution diffusion magnetic resonance imaging (dMRI) since they locate within a ∼1.5mm thin layer of superficial white matter. With the emergence of high-resolution diffusion imaging techniques, this study timely evaluated the effects of image spatial resolution on SAF reconstruction using simulation data and multi-resolution (2, 1.5, and 0.96 mm iso.) empirical data acquired on the same 20 healthy subjects using the gSlider sequence and 20 widely used tractography approaches. Resolution effects were qualitatively assessed through model fitting and tractography results and quantitatively evaluated using global and regional short-range connectivity strength (SCS). It is found that lower resolution systematically reduces SCS across all methods in a spatially varying manner. Moreover, tractography methods exhibit significant differences in resolution sensitivity, with diffusion tensor imaging (DTI) based single-tissue single-fiber tractography showing greater vulnerability than constrained spherical deconvolution (CSD)-based multi-tissue multi-fiber tractography. Probabilistic tracking with anatomical constraints (ACT) and filtering (SIFT) improves robustness. Finally, up-sampling data to nominally higher resolution partially mitigates resolution-induced degradation and improves SAF reconstruction accuracy, particularly for DTI tractography. Based on these findings, higher resolution and multi-shell imaging is recommended if possible. For a given dataset, data up-sampling and DTI-based probabilistic tracking with ACT is recommended for single-shell low b-value data. CSD-based probabilistic tracking with ACT and SIFT is recommended for single-shell higher b-value data and multi-shell data. In summary, this study systematically and quantitatively evaluated resolution effects on SAF reconstruction and structural connectivity estimation and provided practical guidelines for more accurate mapping of SAFs. These advances hold promises to improve the characterization of healthy and diseased human brains in a wide range of neuroscientific and clinical applications.

## 1 Introduction

Short-range association fibers (SAFs) are white matter (WM) pathways that connect adjacent gyri within the same hemisphere (Guevara et al., 2020). These fibers typically span between 3 mm and 30 mm in length and are primarily located within a ∼1.5 mm thin layer of superficial WM beneath the cerebral cortex (Kirilina et al., 2020; Movahedian Attar et al., 2020; Shastin et al., 2022). Due to their characteristic shape following cortical gyration, SAFs are commonly referred to as U-fibers (Tian et al., 2025; Van Dyken et al., 2024).

SAFs are important for understanding the cognitive function and developmental and neurodegenerative mechanisms of human brain. First, they occupy ∼240 cm^3^ of the total ∼420 cm^3^ WM volume and constitute 90% of brain’s axonal fibers (Schüz, 2002), rendering them essential for cortico-cortical communications (Markov et al., 2013). Additionally, due to their latest myelination pattern (Barkovich, 2000; Parazzini et al., 2002; M. Wu et al., 2016) and the “last-in-first-out” principle of neurodegeneration (Braak & Braak, 1996; Fjell et al., 2014), SAFs are among the first structures to show pathological changes in neurodegenerative diseases such as Alzheimer’s (Carmeli et al., 2014; Fornari et al., 2012).

Fiber tracking (tractography) based on diffusion MRI (dMRI) is the most widely used method for *in vivo* mapping of SAFs. DMRI measures the diffusion patterns of water molecules in brain tissue (Le Bihan et al., 1986), which are utilized to infer local microstructure properties (Assaf et al., 2008; Le Bihan et al., 2001; H. Zhang et al., 2012). Tractography integrates local information across voxels to reconstruct continuous fiber pathways and whole-brain connectomes (Mori & Van Zijl, 2002). Pioneering tractography approaches (Mori et al., 1999) relied on principal diffusion direction derived from diffusion tensor imaging (DTI), which are intrinsically limited in resolving crossing fibers. Therefore, more sophisticated methods were developed to model crossing fiber configurations. For instance, the ball-and-stick model separates diffusion signal into isotropic (ball) and anisotropic (stick) components (Behrens et al., 2007; Jbabdi et al., 2012). Similarly, constrained spherical deconvolution (CSD) models each fiber population using an empirically determined response function, or even multiple response functions for different tissue types. (Jeurissen et al., 2014; Tournier et al., 2007).

The accuracy of SAF reconstruction using tractography is hampered by the limited spatial resolution of dMRI. The most widely adopted 2D EPI based dMRI protocol usually employs 2-3 mm iso. spatial resolution and a single b-value of 800-1500 s/mm^2^ for DTI (W. Wu & Miller, 2017) due to the inherent trade-off between the macro- and micro-anatomy resolving capability and signal-to-noise ratio (SNR). These spatial and diffusion resolutions are insufficient for capturing the microstructural patterns of SAFs residing in the ∼1.5mm thin layer of superficial WM (González Ballester et al., 2002), which are even thinner than most cortices. Mixing with gray matter (GM) signals may cause premature termination of tractography or increased uncertainty in probabilistic tracking algorithms, while mixing with deep WM signals may cause SAF tracking to erroneously extend into deeper regions (Reveley et al., 2015), both leading to underestimated SAF connectivity. To map SAFs accurately, both high spatial resolution (1 mm iso. or higher) and high diffusion-encoding sensitivity (2000 s/mm^2^ or higher) are needed to reduce partial volume effects and resolve fiber crossings.

Recent advances in dMRI acquisition have significantly improved imaging resolution. 3D EPI enhances SNR efficiency by shortening TR and enables high isotropic resolution with 3D Fourier encoding, which was first demonstrated useful in *ex vivo* imaging at 0.7 mm iso. resolution (K. L. Miller et al., 2011). For *in vivo* imaging with unavoidable subject motion, 3D multi-slab EPI was developed to mitigate motion artifacts. Each excited slab is corrected for motion-induced shot-to-shot phase variation with a 2D navigator (Engström & Skare, 2013; W. Wu et al., 2016). Recently, this technique was further developed to enable 0.53 mm iso. resolution for *in vivo* dMRI (Z. Li et al., 2024). Another representative technique Generalized Slice Dithered Enhanced Resolution (gSlider) acquires multiple thick slabs using various RF excitations while maintaining high in-plane resolution. These RF-encoded thick slices are then used to compute high-resolution thin slices as an inverse problem with Tikhonov regularization. GSlider improved SNR through multiple excitations and achieved 0.76 mm iso. resolution for *in vivo* dMRI (Setsompop et al., 2018). More recently, Romer-EPTI enabled 0.5 mm iso. resolution for *in vivo* dMRI through super-resolution reconstruction of multiple thick-slice volumes with rotated field-of-view (Dong et al., 2024).

Unfortunately, the benefit of higher image spatial resolution in studying SAFs has not been systematically evaluated despite the emerging high-resolution dMRI techniques. Almost all current SAF studies used conventional low-resolution protocols (K. G. Schilling et al., 2023; Urquia-Osorio et al., 2022; Yuan et al., 2022). The impact of up-sampling on SAF reconstruction is also unclear: while some studies suggested that up-sampling dMRI data prior to modeling and analysis benefits DTI-based tractography (Dyrby et al., 2014), others indicated that data up-sampling may yield results inferior to natively acquired high-resolution data for CSD-based tractography (Dmitri et al., 2019). To bridge these gaps, we quantitatively evaluated the impact of dMRI spatial resolution on reconstructing SAFs and estimating structural connectivity. Our hypothesis is that higher resolution benefits SAF reconstruction through reduced partial volume effects.

Specifically, we comprehensively compared voxel-wise model fitting results, whole-brain fiber tracking results, and structural connectivity estimation results across multiple commonly adopted spatial resolutions. We used both empirical dMRI data acquired with vendor-provided product 2D EPI sequence (1.5 and 2 mm iso.) and the advanced gSlider sequence (0.96 mm iso.) as well as simulation data from the Human Connectome Project (HCP) (Van Essen et al., 2013) (1.25, 1.5, 2, 2.5 mm iso.). We also assessed how fiber models (single or crossing) and tracking methodologies (deterministic or probabilistic, with or without spherical-deconvolutional informed filtering, with or without anatomy constrain) influence across-resolution differences. These findings elucidate the impacts of dMRI spatial resolution on short-range and whole-brain structural connectivity estimation, provide a practical guideline for optimized dMRI acquisition and analysis protocols, and potentially advance our understanding of SAFs and human brain networks and their changes.

## 2 Material and Methods

### 2.1 Empirical Data

With approval from the Tsinghua University Institutional Review Board and written informed consent, MRI data were acquired from twenty healthy young adults (mean age: 23.45 ± 1.80 years; 12 females) on a Siemens MAGNETOM Prisma 3T scanner equipped with a 32-channel head coil. T1-weighted MPRAGE (1 mm iso.) and dMRI data at multiple isotropic resolutions (0.96, 1.5, and 2 mm iso.) were acquired for each subject.

The submillimeter resolution (0.96 mm iso.) diffusion data were acquired using the gSlider sequence (Setsompop et al., 2018). This sequence acquired five 4.8 mm-thick slabs with 0.96×0.96 mm^2^ in-plane resolution using five optimized and varying RF profiles for excitation, which were then used to reconstruct 0.96 mm-thick slices for achieving 0.96 mm iso. spatial resolution. To minimize slab-boundary artifacts, low-resolution B1+ maps were acquired to calibrate the RF encoding profiles for accounting for incomplete T1 recovery effects due to B1+ field inhomogeneity (Liao et al., 2020). The diffusion protocol included 32 diffusion weighted images (DWIs) at b = 1000 s/mm^2^ and 64 DWIs at b = 2500 s/mm^2^ along uniformly distributed diffusion directions. A b = 0 image was inserted after every 16 DWIs. An additional b = 0 image with reversed phase-encoding direction was acquired for susceptibility distortion correction.

Diffusion data at commonly adopted 1.5 mm and 2 mm iso. resolution was acquired using the product 2D simultaneous multi-slice (SMS) pulsed gradient spin echo (PGSE) single-shot EPI sequence (i.e., ep2d sequence). The diffusion-encoding scheme (b-values and directions) and phase-encoding strategy were identical to those used in the gSlider sequence. Detailed acquisition parameters are provided in Table 1.

**Table 1.**
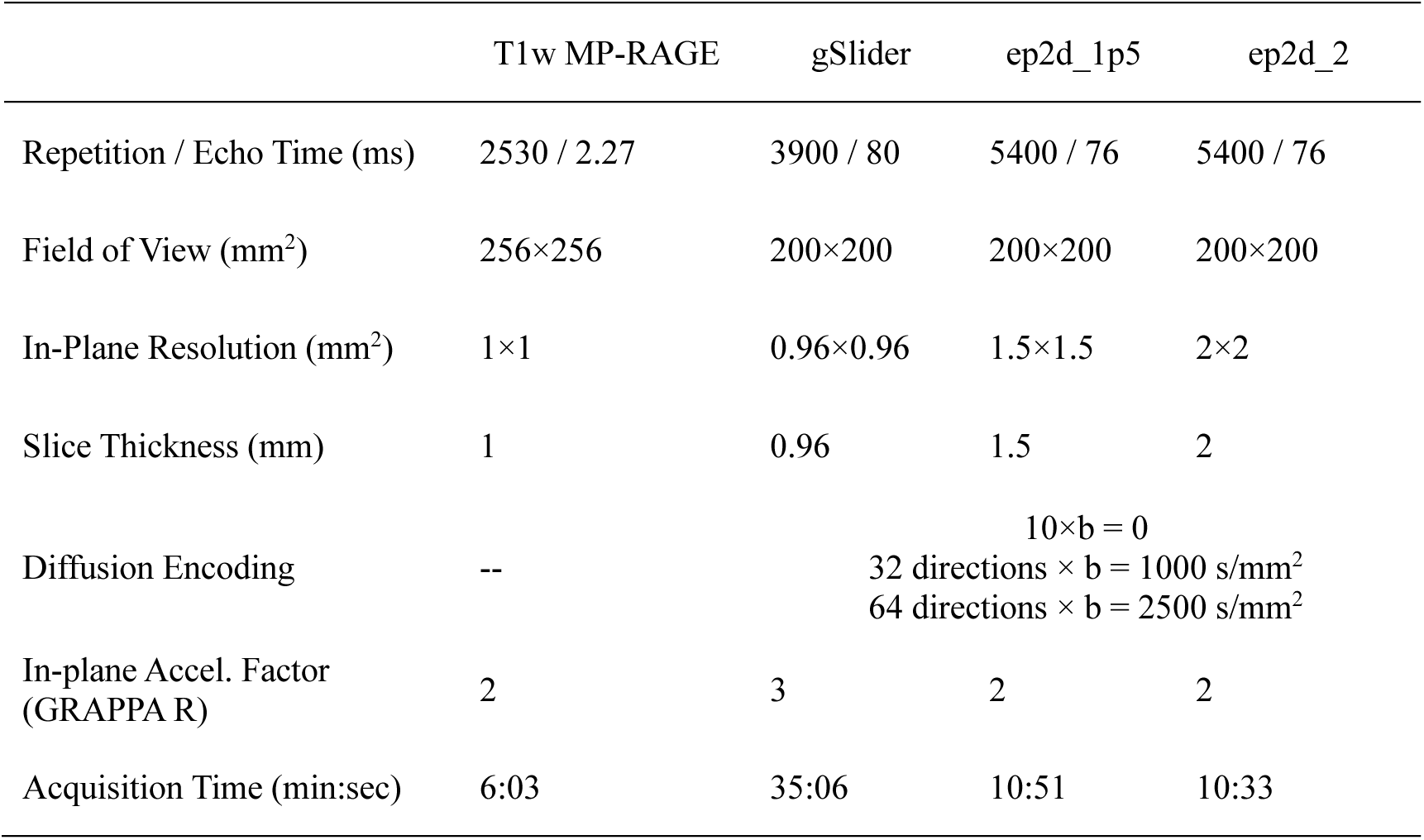
Acquisition parameters for empirical data.

All diffusion MRI data were preprocessed for distortion correction and image alignment. First, gradient nonlinearity correction was applied (Janke et al., 2004). Then, the susceptibility-induced off-resonance field was estimated using b = 0 images acquired with opposite phase encoding directions by the “topup” function from the FMRIB Software Library (FSL) (Andersson et al., 2003; Graham et al., 2017). This field map, along with the diffusion data, were input to FSL’s “eddy” function for correcting susceptibility and eddy current induced distortions and subject motion (Andersson et al., 2016). Finally, to investigate the effects of up-sampling (Dyrby et al., 2014), the preprocessed data at 1.5 and 2 mm iso. resolution were up-sampled to 0.96 mm iso. using FSL’s “flirt” function with spline interpolation.

T1w images were also corrected for gradient nonlinearity. Cortical surface reconstruction and volumetric segmentation were performed using FreeSurfer’s “recon-all” function (Fischl, 2012). The preprocessed diffusion data and T1w data were co-registered using the boundary-based registration implemented in FreeSurfer’s “bbregister” function (Greve & Fischl, 2009).

### 2.2 Simulation Data

To complement empirical data analysis, simulation experiments were conducted using pre-processed and co-registered T1w (0.7 mm iso.) and dMRI (1.25 mm iso.) data from 20 healthy subjects from the HCP WU-Minn consortium (Van Essen et al., 2013). The dMRI protocol included 18 b = 0 volumes and 90 DWIs for each of the three shells (b = 1000, 2000, 3000 s/mm²), with acquisition and preprocessing details previously described (Glasser et al., 2013; Sotiropoulos et al., 2013; Uğurbil et al., 2013).

To simulate the dMRI data at multiple spatial resolutions, the original dMRI data at 1.25 mm iso. were down-sampled to 1.5, 2, and 2.5 mm iso. resolution. The down-sampling was performed by applying a Hanning window to the frequency space of the 3D image data to set high-frequency components beyond the cut-off frequency to zero while minimize Gibbs ringing artifacts. The nominal resolution of the down-sampled data was still 1.25 mm.

### 2.3 Connectivity quantification

Multiple tractography pipelines from the MRtrix3 software were adopted for estimating SAF pathways and structural connectivity (Figure 1). Specifically, the tractography was performed on single-shell or multi-shell data using: (1) single-fiber (tensor) or crossing-fiber (constrained spherical deconvolution (CSD)) modeling and (2) deterministic or probabilistic tracking, (3) with or without Spherical-deconvolution Informed Filtering of Tractograms (SIFT), (4) with or without anatomical constrain tracking (ACT). SIFT and ACT improve the biological plausibility of reconstructed fiber pathways by selectively removing streamlines, making the streamline density along each fiber direction proportional to the corresponding fiber orientation distribution (FOD) amplitude (Smith et al., 2013, 2015) and forcing them to initiate and terminate at the interface of WM and cortical or deep GM (Smith et al., 2012), respectively. In total, 20 tractography strategies were used (Table 2), with their detailed tracking parameters provided in Supplementary Information Table S1 (empirical data) and Table S2 (simulation data). For the single-shell data, data with b = 1000 s/mm^2^ or b = 2500 s/mm^2^ from empirical data and data with b = 1000 s/mm^2^ or b = 2000 s/mm^2^ from simulation data was used. For the multi-shell data, data with b = 1000 s/mm^2^ and b = 2500 s/mm^2^ from empirical data and data with b = 1000 s/mm^2^, b = 2000 s/mm^2^ and b = 3000 s/mm^2^ from simulation data were used.

**Figure 1.**
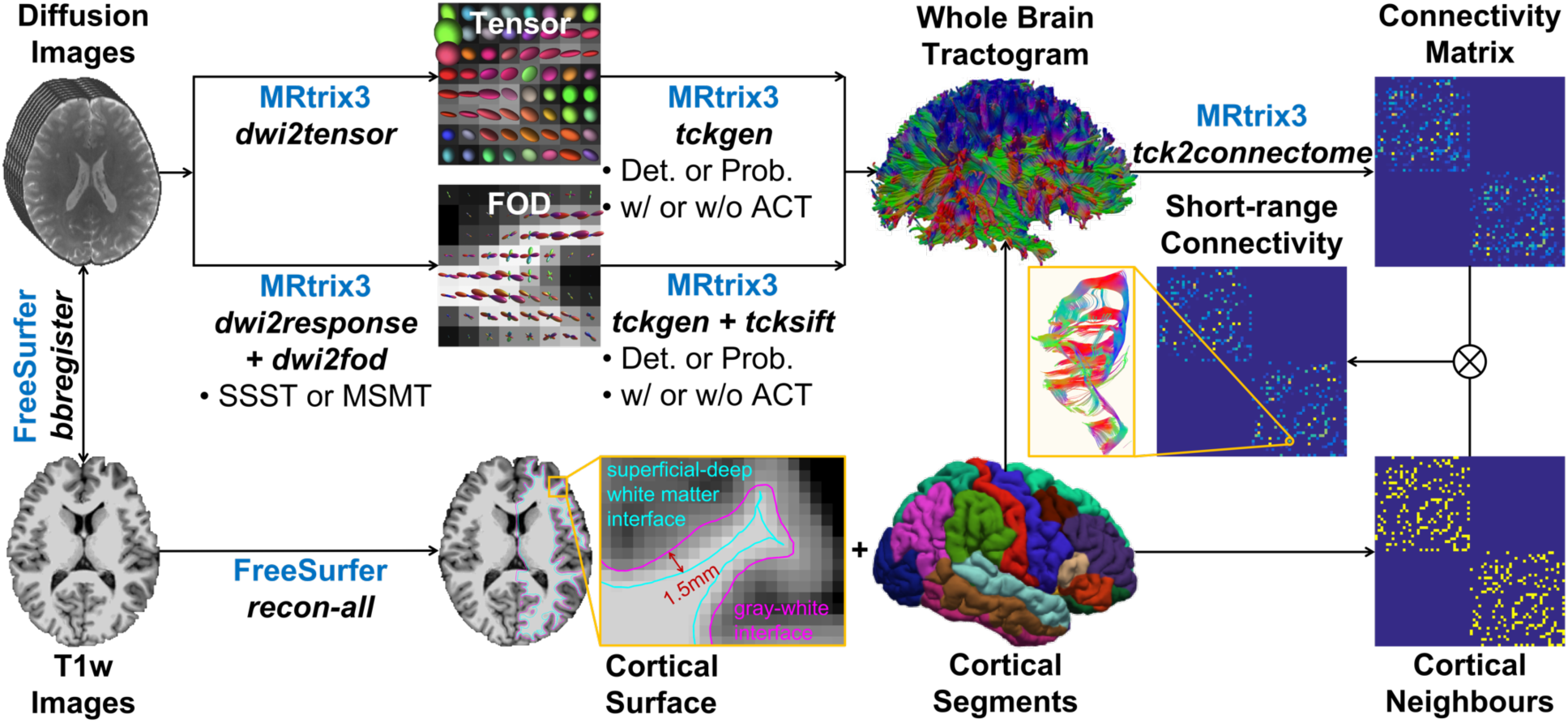
Structural connectivity estimation overview. Tractography was performed using the MRtrix3 software on single-shell or multi-shell empirical and simulation diffusion data using: (1) single-fiber (tensor modeling) or crossing-fiber modeling (single-shell single-tissue (SSST) or multi-shell multi-tissue (MSMT) constrained spherical deconvolution (CSD)) and (2) deterministic or probabilistic tracking, (3) with or without Spherical-deconvolution Informed Filtering of Tractograms (SIFT), (4) with or without Anatomical Constrain Tracking (ACT) for estimating structural connectivity assisted by the FreeSurfer software.

**Table 2.** Overview of tractography strategies.

| Single-shell data |  |  |  |
| --- | --- | --- | --- |
|  | Single-fiber model w/o SIFT | Crossing-fiber model w/o SIFT | Crossing-fiber model w/ SIFT |
| Det. | Low B, Sing, Det, w/o SIFT, w/o ACT<br>Low B, Sing, Det, w/o SIFT, w/ ACT | High B, Xing, Det, w/o SIFT, w/o ACT<br>High B, Xing, Det, w/o SIFT, w/ ACT | High B, Xing, Det, w/ SIFT, w/o ACT<br>High B, Xing, Det, w/ SIFT, w/ ACT |
| Prob. | Low B, Sing, Prob, w/o SIFT, w/o ACT<br>Low B, Sing, Prob, w/o SIFT, w/ ACT | High B, Xing, Prob, w/o SIFT, w/o ACT<br>High B, Xing, Prob, w/o SIFT, w/ ACT | High B, Xing, Prob, w/ SIFT, w/o ACT<br>High B, Xing, Prob, w/ SIFT, w/ ACT |

| Multi-shell data |  |  |
| --- | --- | --- |
|  | Crossing-fiber model w/o SIFT | Crossing-fiber mode w/ SIFT |
| Det. | Multi B, Xing, Det, w/o SIFT, w/o ACT<br>Multi B, Xing, Det, w/o SIFT, w/ ACT | Multi B, Xing, Det, w/ SIFT, w/o ACT<br>Multi B, Xing, Det w/ SIFT, w/ ACT |
| Prob. | Multi B, Xing, Prob, w/o SIFT, w/o ACT<br>Multi B, Xing, Prob, w/o SIFT, w/ ACT | Multi B, Xing, Prob, w/ SIFT, w/o ACT<br>Multi B, Xing, Prob, w/ SIFT, w/ ACT |

Abbreviations: Low B = single-shell with low b-value; High B = single-shell with high b-value; Multi B = multiple shells with both low and high b-values; Sing = single-fiber model; Xing = crossing-fiber model; Det=deterministic; Prob=probabilistic; SIFT = Spherical-deconvolution Informed Filtering of Tractograms; ACT = anatomical constraints tractography.

The tensor-based tractography was performed on single-shell data with a low b-value (b = 1000 s/mm^2^) using the “tckgen” function from the MRtrix3 software (Tournier et al., 2019) using both deterministic (“Tensor_Det” option) and probabilistic (“Tensor_Prob” option) tracking, without SIFT, and with or without ACT.

The CSD was performed with “dwi2response” and “dwi2fod” functions from the MRtrix3 software (Jeurissen et al., 2014; Tournier et al., 2004, 2007) using single-shell single-tissue (SSST) or multi-shell multi-tissue (MSMT) approaches. SSST-CSD utilized single-shell data with a high b-value (b = 2500 s/mm^2^ for empirical data and b = 2000 s/mm^2^ for simulation data) and MSMT-CSD utilized multi-shell data. Subsequently, CSD-based tractography was performed using the “tckgen” function using both deterministic (“SD_Stream” option) and probabilistic (“iFOD2” option) tracking, with or without SIFT (applied with “tcksift” function in MRtrix3 software) and with or without ACT.

Structural connectivity matrix 𝑊 was generated using the 68 cortical regions defined by the Desikan-Killiany (DK) atlas (Desikan et al., 2006). The element 𝜔*_i,j_* of the matrix 𝑊, representing the connection strength between region 𝑖 and region 𝑗, was defined as:

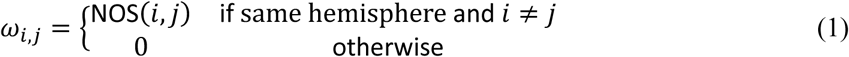

where NOS(𝑖, 𝑗) is the number of streamlines connecting region 𝑖 and region 𝑗. 𝑊 includes structural connectivity of association fibers but not projection and commissural fibers.

### 2.4 Neighboring Cortices Identification

To identify short-range connectivity, the neighboring cortices of each cortical region was identified. The neighboring pattern was represented by a matrix 𝐴(𝑖, 𝑗), with 𝐴(𝑖, 𝑗) = 1 if a cortical region 𝑖 from the DK atlas borders on cortical region 𝑗. Otherwise, 𝐴(𝑖, 𝑗) = 0. Therefore, the connectivity strength 𝜔*_i,j_* at location with 𝐴(𝑖, 𝑗) = 1 in the connectivity matrix 𝑊 results from SAFs, while 𝜔_i,j_ at location with 𝐴(𝑖, 𝑗) = 0 results from long-range association fibers.

### 2.5 Connectivity Metrics

Two quantitative metrics were derived from the connectivity matrix 𝑊 to evaluate the effect of spatial resolution on SAF reconstruction, including global short-range connectivity strength (GSCS), regional SCS (RSCS). Specifically, GSCS was defined as the ratio of the short-range connectivity strength to the entire connectivity strength:

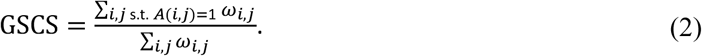

For a particular cortical region 𝑘, the regional SCS RSCS*_k_* was calculated as the ratio of its short-range connectivity strength to the entire connectivity strength originating from this region:

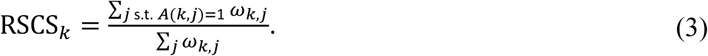

## 3. Results

The empirical diffusion data and HCP simulation data at different spatial resolutions and derived microstructural metrics exhibited high quality, with adequate signal-to-noise ratio (SNR) and no noticeable artifacts (Figure 2, Supplementary Information Figure S1). Structural details (e.g., GM-WM interface) were clearer while SNR of DTI metric maps were slightly lower at higher resolution, as expected.

**Figure 2.**
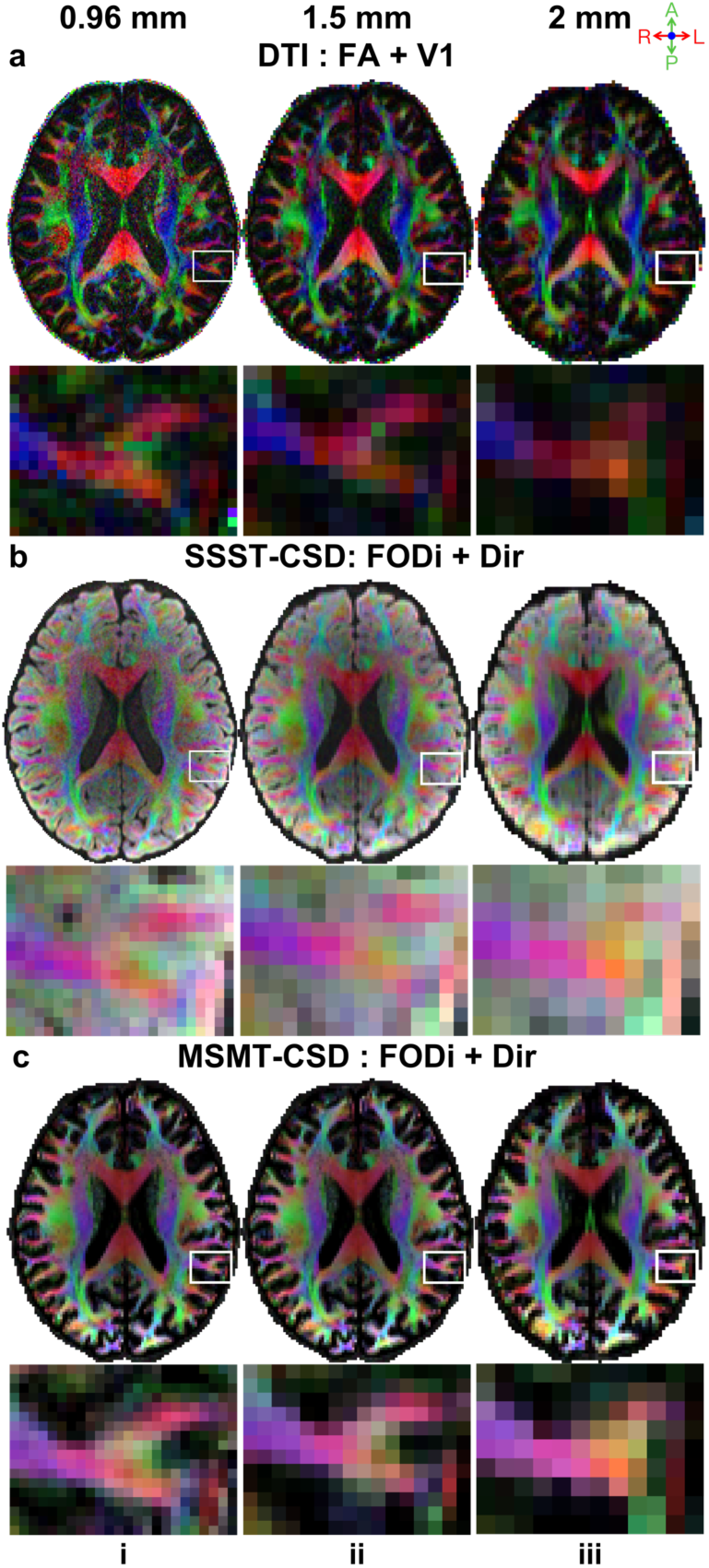
Empirical data quality. Axial slices from a representative subject with fitted results from three diffusion models (a: DTI, b: SSST-CSD, c: MSMT-CSD) across three resolutions (i: 0.96 mm, ii: 1.5 mm, iii: 2 mm) are displayed. The fractional anisotropy (FA) map color-coded by primary vector (V1) (a) and the fiber orientation distribution integral (FODi) map color-coded by the overall fiber direction (b, c) are shown for DTI and CSD methods (red: left–right; green: anterior–posterior; blue: superior–inferior), respectively.

Higher spatial resolution substantially improved the representation of superficial WM architecture (Figure 3). At 0.96 mm iso. resolution, SAFs were clearly delineated by the DTI primary eigenvectors (i.e., primary fiber orientations) (Figure 3a, ii blue arrows, green fibers). They diminished as the spatial resolution decreased (Figure 3a, iii, v) and entirely disappeared at 2 mm iso. resolution (Figure 3a, v), in which case the superficial WM layer was visually dominated by reddish long-range fibers entering the cortex. Up-sampling the 1.5 mm iso. data increased the number of voxels representing SAFs and thus slightly improved the SAF delineation (Figure 3a, iii to Figure 3a, iv). Up-sampling the 2 mm iso. diffusion data did not recover any SAFs that were originally missing at the native resolution (Figure 3a, v to Figure 3a, vi). Superficial WM layer FA was higher at 0.96 mm iso. resolution (Figure 3a, ii) and decreased at lower resolutions (Figure 3a, ii–vi) due to the partial volume effect with GM and the radial long-range fibers entering the cortex.

**Figure 3.**
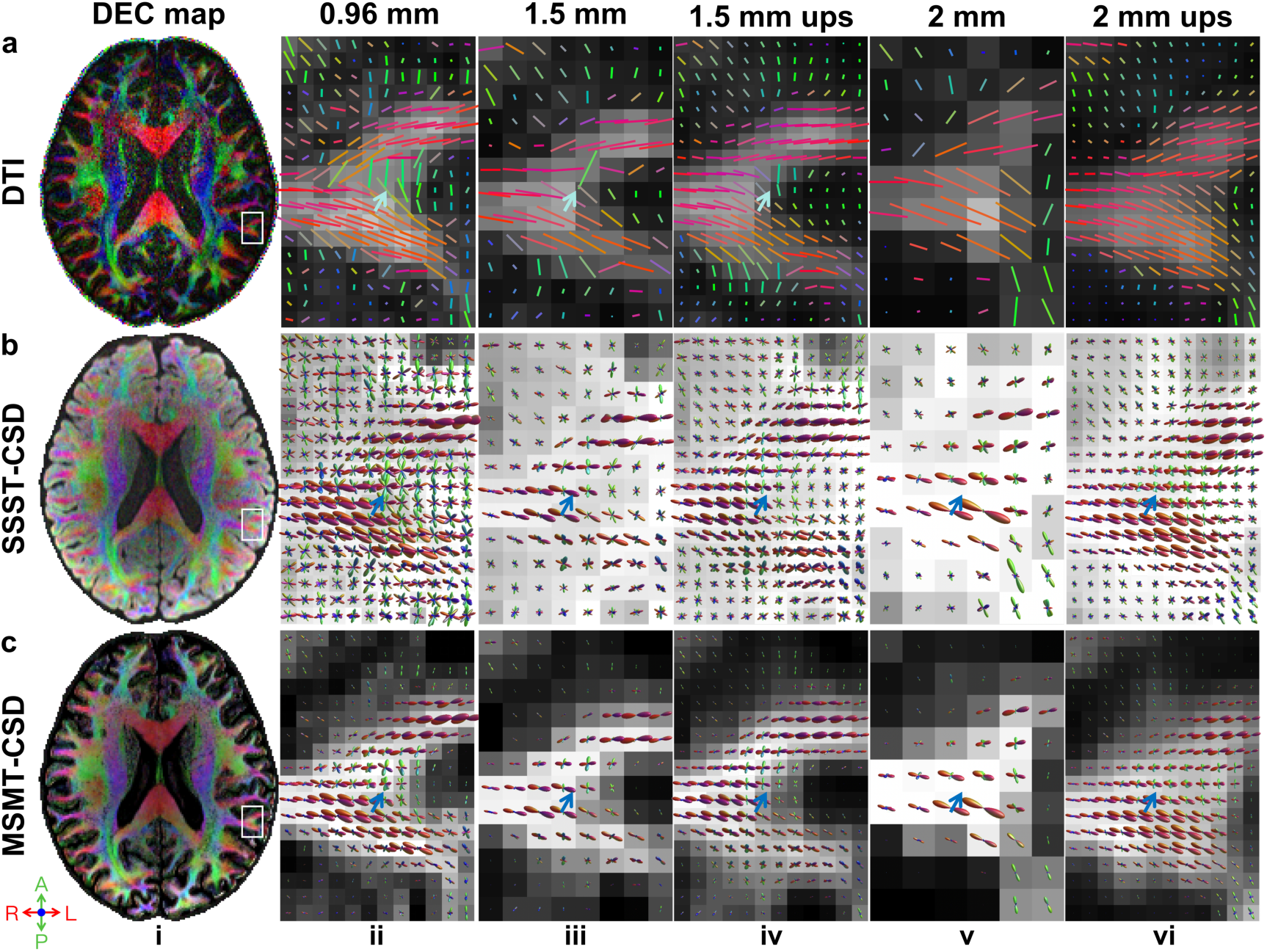
Axon orientations from empirical data. Primary fiber direction encoded color (DEC) maps (red: left– right; green: anterior–posterior; blue: superior–inferior) at 0.96 mm iso. resolution from a representative subject (i) are displayed for three methods including DTI (a, i), SSST-CSD (b, i), and MSMT-CSD (c, i), with a region of interest (white boxes) containing a gyrus, subcortical white matter, and their interface shown in enlarged views overlaid on FA (a), and FOD integral (b, c) maps across three different native spatial resolutions (ii: 0.96 mm, iii: 1.5 mm, v: 2 mm) and two nominally high 0.96 mm iso. resolution up-sampled from 1.5 mm and 2 mm iso. resolution (iv: 1.5 mm up-sampled, vi: 2 mm up-sampled). Blue arrows highlight a region with short-range association fibers.

SAFs were robustly delineated across spatial resolutions by SSST-CSD and MSMT-CSD (Figures 3b-c), crossing-fiber models that are capable of disentangling SAFs from long-range fibers entering the cortex. However, the FOD magnitude (reflecting the volume fraction) of SAFs gradually reduced as the resolution decreased due to the partial volume effect, potentially hampering the accurate tracking of SAFs and the faithful estimation of short-range connectivity. It is worth noting that the commonly adopted SIFT method filters track density according to the FOD magnitude. Up-sampling the lower resolution data did not recover the lost FOD magnitude of SAFs (Figure 3b-c, iv and vi). Comparing to SSST-CSD results, MSMT-CSD results included much fewer fibers with lower FOD magnitude in the superficial WM (i.e., mainly SAFs) and cortex (Figure 3c, blur arrows), which were potentially spurious. Moreover, the FOD integral map from MSMT-CSD exhibited clear GM-WM boundaries (Figure 3c), indicating this model’s capability to distinguish between GM and WM, which were not displayed in SSST-CSD maps. Similar voxel-wise fiber patterns for DTI, SSST-CSD, and MSMT-CSD were observed in simulation data (Supplementary Information Figure S2).

U-shaped SAFs were reliably reconstructed by different tractography methods, as confirmed by visual inspection, e.g., SAFs connecting the right middle and superior temporal regions (Figure 4, Supplementary Information Figure S3). The track density reduced as the spatial resolution decreased. Up-sampling the diffusion data slightly increased the tract density (Fig. 4ii vs. iii, iv vs. v). Track pathways from DTI and MSMT-CSD were more coherent comparing to those from SSST-CSD, which possibly contained spurious short fibers at the bottom of the entire tract.

**Figure 4.**
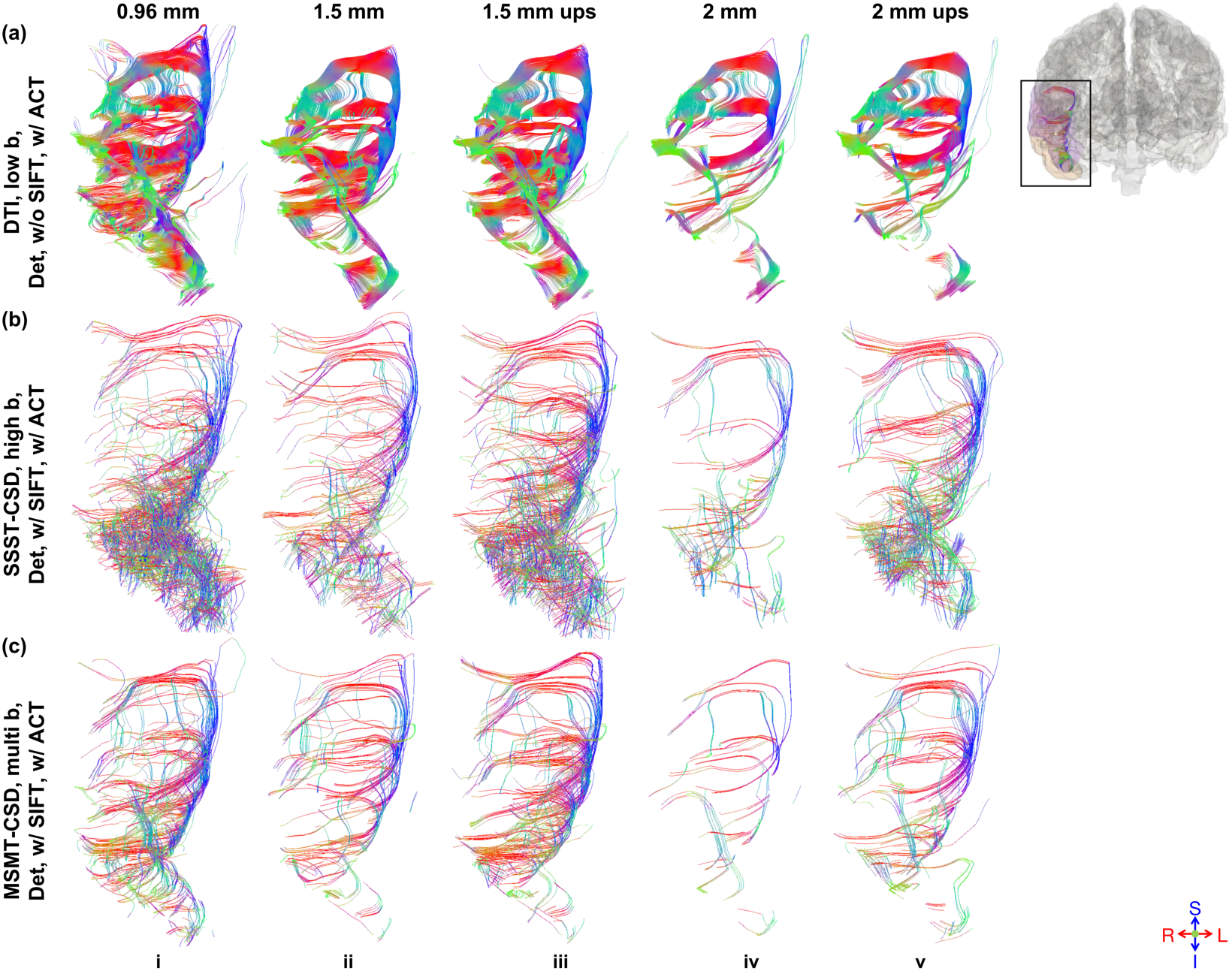
Empirical data tractography results. U-shaped SAFs connecting the right middle and superior temporal gyri (black box) reconstructed using three modeling methods (a: DTI on low b-value single-shell data, b: SSST-CSD on high b-value single-shell data with SIFT, c: MSMT-CSD on multi b-value data with SIFT) and deterministic tractography with ACT across three different native spatial resolutions (i: 0.96 mm, ii: 1.5 mm, iv: 2 mm) and two nominally high 0.96 mm iso. resolution up-sampled from 1.5 and 2 mm iso. resolution (iii: 1.5 mm up-sampled, v: 2 mm up-sampled) are displayed.

Lower spatial resolution consistently led to decreased GSCS (Figure 5, Supplementary Information Figure S4). The magnitude of this resolution induced GSCS reduction varied across modeling and tracking methods. Regarding fiber modeling methods, DTI single-fiber models exhibited the largest decreases in GSCS (Fig. 5a, i-iv), while CSD-based crossing fiber models showed lower changes (Figure 5a, v-xii and 5b).

**Figure 5.**
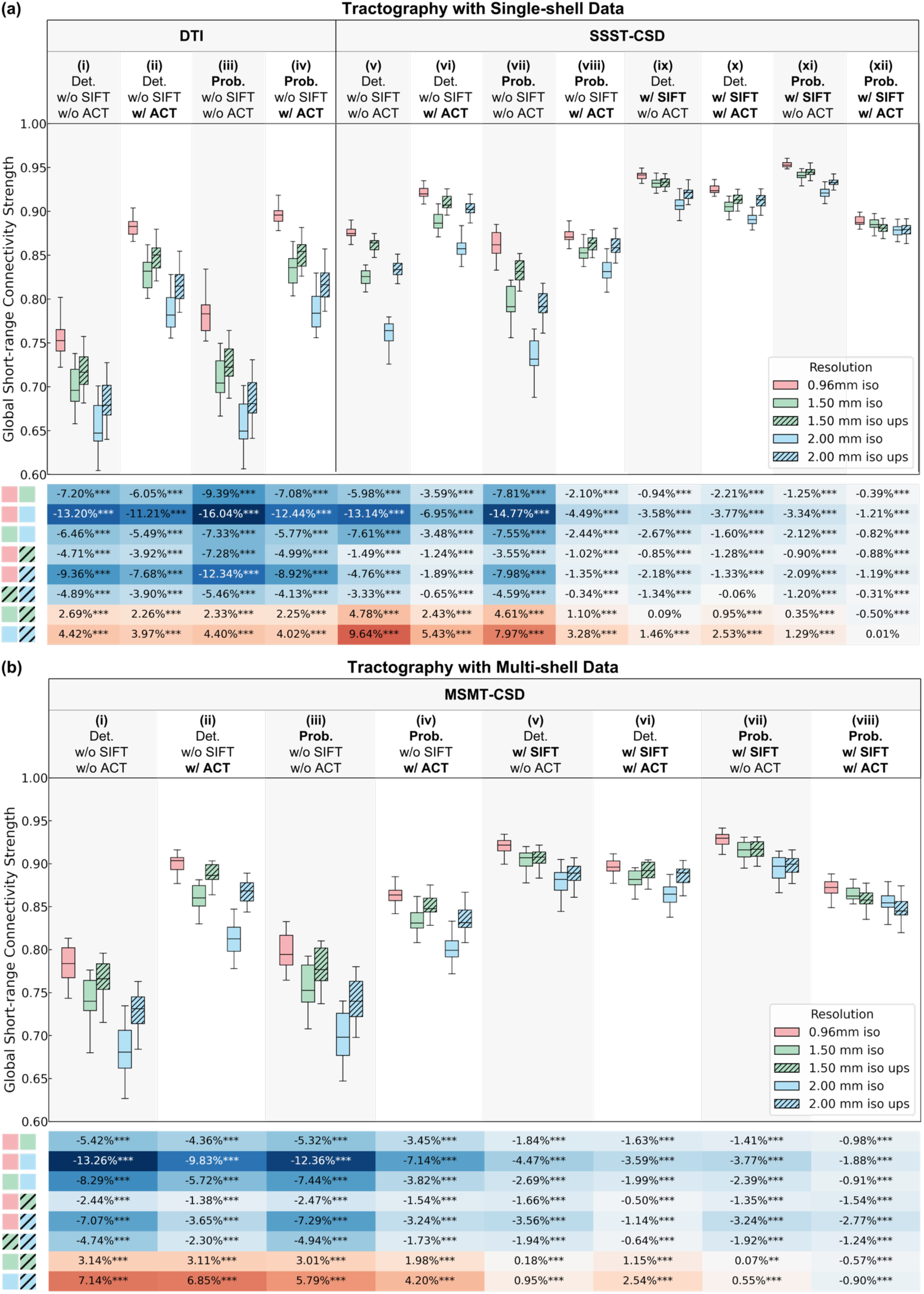
Empirical data GSCS. Box plots for GSCS from different tractography methods using single-shell (a) and multi-shell (b) data at three native spatial resolutions (red: 0.96 mm iso., green: 1.5 mm iso., blue: 2 mm iso.) and two nominally high 0.96 mm iso. resolution up-sampled from 1.5 mm and 2 mm iso. resolution (hatched green: 1.5 mm up-sampled, hatched blue: 2 mm up-sampled) display the distribution (i.e., median, interquartile range, and range) of GSCS from 20 subjects in the upper panel. The lower table lists the relative difference of GSCS at lower resolution compared to the GSCS at higher resolution (each row) for different tractography methods (each column), with asterisks denoting significance levels (*: p < 0.05, **: p < 0.01, ***: p < 0.001). The color of each table cell indicates the magnitude and direction of the GSCS difference (blue: decrease, red: increase)

Regarding tracking methods, probabilistic and deterministic algorithms exhibited comparable sensitivity to resolution changes for single-fiber model (Figure 5a, iii, iv vs. Figure 5a, i, ii), while deterministic algorithms exhibited higher sensitivity for crossing-fiber models (Figure 5a, v, vi, ix, x and Figure 5b, i, ii, v, vi vs. Figure 5a, vii, viii, xi, xii and 5b, iii, iv, vii, viii). Moreover, GSCS obtained using SIFT (Figure 5a, ix-xii and 5b, v-viii) showed greater robustness to resolution changes than without using SIFT (Figure 5a, v-viii and 5b, i-iv). Finally, methods lacking anatomical constraints (Figure 5a, i, iii, v, vii, ix, xi and 5b, i, iii, v, vii) consistently showed larger GSCS reductions compared to methods incorporating ACT (Figure 5a, ii, iv, vi, viii, x, xii and 5b, ii, iv, vi, viii).

Data up-sampling generally increased GSCS, partially offsetting the reduction caused by lower native resolution across most methods. However, for CSD-based probabilistic tractography incorporating both SIFT and ACT, the mostly adopted tracking option, up-sampling data might even further reduce GSCS (Figure 5a, xii and 5b, viii). Generally, tracking methods without using SIFT or ACT benefitted more from data up-sampling.

Lower spatial resolution led to decreased RSCS for most cortical regions, while the effect and spatial pattern varied across tractography methods (Figure 6). DTI-based tractography was the most sensitive to spatial resolution reduction. Its derived RSCS substantially decreased for almost all cortical regions at lower resolution, particularly the superior temporal and precuneus cortex (Figure 6a). For SSST-CSD-based tractography, RSCS of 37 cortical regions significantly decreased at lower resolution (Figure 6b), with the pars opercularis and isthmus cingulate cortex exhibiting the most pronounced effects. RSCS of 18 cortical regions slightly increased. For MSMT-CSD-based tractography, most cortices exhibited RSCS reduction while 4 cortices including left cuneus, right cuneus, right lateral occipital and right postcentral exhibited RSCS increase at lower resolution (Figure 7c). The pars opercularis, and para hippocampal cortex demonstrated the highest resolution sensitivity. All spatial heterogeneity results for different methods and resolution comparisons (i.e., native 1.5 mm iso., 2 mm iso., and up-sampled nominal 0.96 mm iso. vs. 0.96 mm iso) and simulation data are provided in Supplementary Information Figures S5 (empirical) and S6 (simulation).

**Figure 6.**
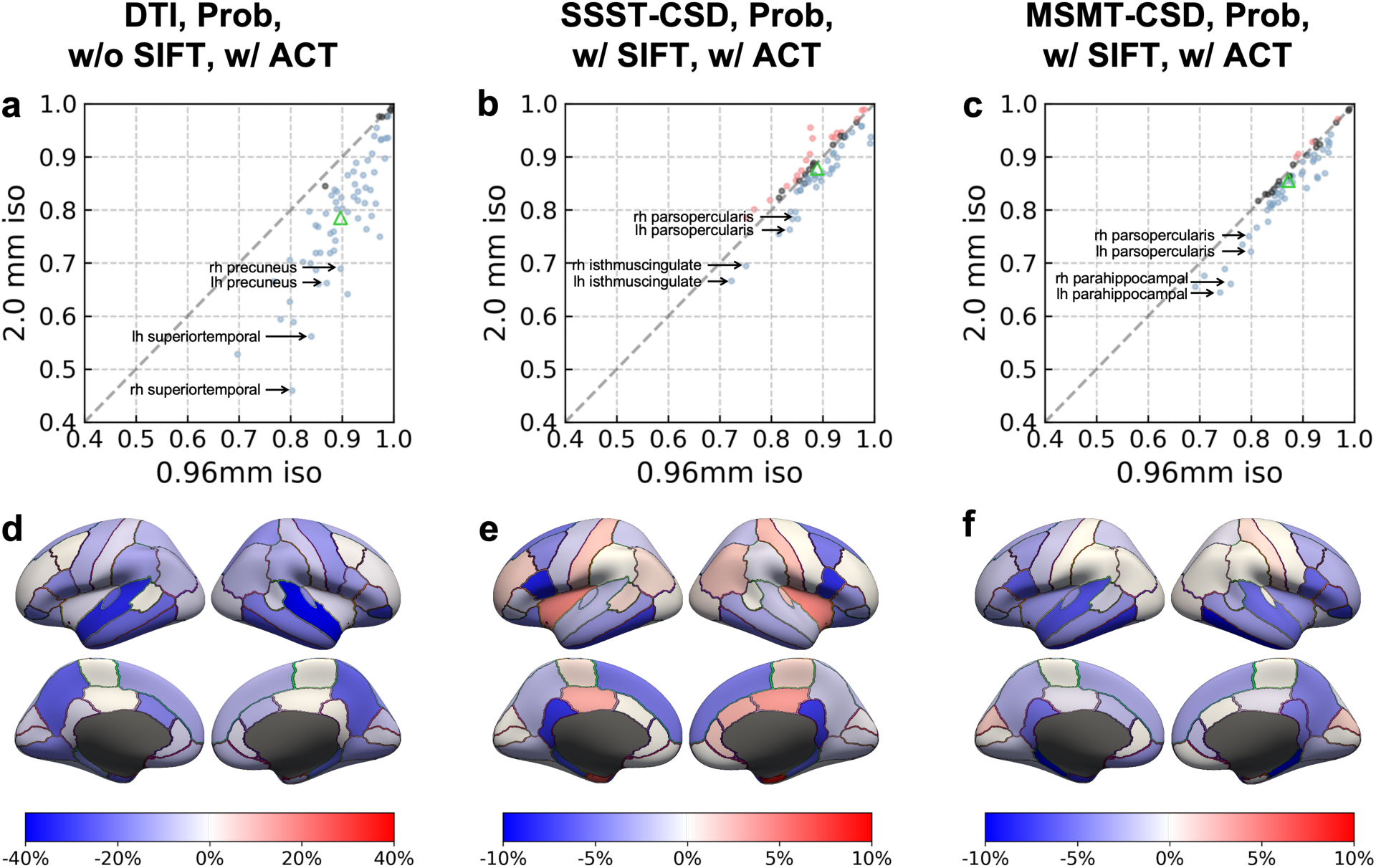
Empirical data RSCS. Scatter plots comparing RSCS at high resolution (0.96 mm iso., x-axis) versus at low resolution (native 2 mm iso., y-axis) are displayed with each point representing a cortical region, color representing the difference and its significance (blue = decrease, red = increase, black = none), and the green triangle representing GSCS for anatomically constrained probabilistic tractography with different modeling and tracking options (a: DTI on low b-value single-shell data without SIFT, b: SSST-CSD on high b-value single-shell data with SIFT, c: MSMT-CSD on multi b-value data with SIFT). The difference of RSCS at low resolution compared to RSCS at high resolution for each cortical region is displayed on inflated surfaces, with color and its intensity representing the pattern.

## 4. Discussion

Our study systematically quantified the resolution impact on SAF reconstruction by comparing model fitting results, tractography outputs, and connectivity-based quantitative metrics. Degradation of the superficial WM architecture at lower resolution was observed in model fitting results, primarily attributable to increased partial volume effects. Tractography results indicated the presence of well-formed U-shaped fibers across all resolutions, but with significantly lower fiber density at lower resolutions. Quantitative analyses further revealed that lower resolution leads to an underestimation of the proportion of SAF within the connectome. Based on these findings, the adoption of high-resolution dMRI for SAF research is recommended whenever feasible, and specific post-processing recommendations are provided for scenarios where high-resolution acquisition is constrained.

Our study systematically quantified the effects of dMRI spatial resolution and tractography methodology on SAF reconstruction and structural connectivity estimation. While the neuroimaging community generally accepts that higher spatial resolution improves the visualization of SAFs and often demonstrated this phenomena qualitatively in studies showcasing new high resolution dMRI sequences (Dong et al., 2024; Z. Li et al., 2024; Setsompop et al., 2018), systematic and quantitative supporting evidence has been lacking. Previous across-resolution comparisons suffered from several limitations, including the reliance on qualitative assessments without quantification (Song et al., 2014), the use of indirect comparisons across different datasets or cohorts that hinders direct interpretation (F. Zhang et al., 2024), or a restricted scope only evaluating a single tractography pipeline (Reveley et al., 2015; Song et al., 2014; F. Zhang et al., 2024). Our study confronts these gaps by providing quantitative assessments using connectivity-based metrics of GSCS, RSCS and subsequent network topological properties such as small-worldness (details in Supplementary Information) (Jiang et al., 2021) and direct comparisons across 3 spatial resolutions and 20 distinct tractography pipelines, comprehensively evaluating critical factors including spatial resolution, data up-sampling, diffusion-encoding sensitivity, deterministic or probabilistic tractography, SIFT and ACT, on both simulation and empirical multi-resolution data. Thanks to the state-of-the-art high-resolution dMRI sequence gSlider, the 0.96 mm iso. dMRI data and the 1.5 and 2 mm iso. data were acquired on the same 20 healthy individuals, providing a direct across-resolution, quantified and systematic comparison on empirical data for the first time. This comprehensive approach establishes a rigorous base for optimizing dMRI acquisition and processing strategies.

The fundamental reason that hampers the accurate reconstruction of SAFs at lower resolutions is the signal partial volume effect that degrades voxel-wise fiber modeling and delineation (Figure 3 and Supplementary Information Figure S2). It primarily occurs through a complex inter-tissue and inter-fiber mixing of signals from GM, SAFs (tangential to the cortical fold) and long-range fibers entering the cortex (radial to the cortical fold) at the superficial WM layer. Generally, GM signal contamination reduces the FA or FOD integral, potentially causing premature streamline termination. Long-range fiber contamination increases the volume fraction of radial fibers, potentially reducing the representation of SAFs and increasing the possibility for erroneous fiber pathways, a well-studied “crossing-fiber” problem in dMRI community (K. Schilling et al., 2017). Single-tissue single-fiber method like DTI (Fig. 5a, -7.4% GSCS change from 0.96 to 1.5 mm iso., -13.1% GSCS change from 0.96 to 2 mm iso.,) is more prone to the signal partial volume effect and is therefore less robust to image resolution decrease compared to the multi-fiber method like SSST-CSD (Fig. 5a, -3.0 % GSCS change from 0.96 to 1.5 mm iso., -6.3 % GSCS change from 0.96 to 2 mm iso.) and MSMT-CSD (Fig. 5b, -3.0 % GSCS change from 0.96 to 1.5 mm iso., -6.9 % GSCS change from 0.96 to 2 mm iso.). This suggests that resolution-dependent effects should be taken into consideration when comparing network metrics across studies and centers.

The choice of tracking strategies significantly influences the sensitivity of SAF estimation to dMRI spatial resolution. Probabilistic tracking showed greater robustness to resolution decrease (Fig. 5, -3.8% GSCS change from 0.96 to 1.5 mm iso., -7.5% GSCS change from 0.96 to 2 mm iso.) compared to deterministic tracking (Fig. 5, -3.8% GSCS change from 0.96 to 1.5 mm iso., -8.1% GSCS change from 0.96 to 2 mm iso.). Tractography without ACT and SIFT resulted in lower GSCS robustness across resolution (Fig. 5, - 6.2% GSCS change from 0.96 to 1.5 mm iso., -13.4% GSCS change from 0.96 to 2 mm iso.). Unconstrained seeding (uniform across the brain volume) places more seeds along long-range fibers than along SAFs, leading to disproportionate long-range fibers reconstruction that overshadows SAFs. ACT strategy mitigates this problem by restricting seeding to the GM-WM interface (Smith et al., 2012), while SIFT refines tractograms by matching streamline density to FOD amplitude, removing redundant long-range fibers (Smith et al., 2013). Consequently, combining ACT and SIFT yielded the most robust GSCS across resolution and likely the most biologically accurate SAF reconstruction and connectome estimation (Fig. 5, -1.3% GSCS change from 0.96 to 1.5 mm iso., -2.6% GSCS change from 0.96 to 2 mm iso.).

Up-sampling low-resolution data generally restores some lost GSCS, with exceptions. For most tractography pipelines, data up-sampling increased GSCS to the level higher than native low-resolution data GSCS but lower than native high-resolution data GSCS. This effect was particularly pronounced for DTI-based methods, suggesting that up-sampling can partially mitigate partial volume effects and reveal finer anatomical details in DTI, consistent with previous findings (Dyrby et al., 2014). For CSD-based methods, while the estimated FODs accurately represent the signal within each voxel, the reduced proportion of tangential fibers at lower resolutions poses challenges for tractography. Data up-sampling alleviated some of these difficulties, yet it typically failed to restore GSCS to native high-resolution levels (Dmitri et al., 2019). Notably, for the recommended and mostly widely used CSD probabilistic tractography equipped with both SIFT and ACT strategies, the initial fiber tracking was already robust (Fig. 5, -0.7% GSCS change from 0.96 to 1.5 mm iso., -1.5% GSCS change from 0.96 to 2 mm iso.), data up-sampling offered minimal additional improvement in GSCS. This finding suggests that acquisition spatial resolution might be another critical factor that requires careful considerations when processing multi-center dMRI data (i.e., dMRI data harmonization (L. Ning et al., 2020)) in addition to diffusion-encoding sensitivity and direction.

Based on our findings, we provide practical guidelines for optimized dMRI strategies for SAF reconstruction and characterization. First, high-resolution (preferably sub-millimeter) dMRI is recommended if possible given adequate SNR and a reasonable scan time. Our study demonstrated that sequences like gSlider achieved this within a practical timeframe (∼30 minutes) for research, while alternative approaches including 3D multi-slab EPI (Engström & Skare, 2013), MB-MUSE (Bruce et al., 2017; Chen et al., 2013), SMSlab (Liu et al., 2023), EPTI (Wang et al., 2019), Romer-EPTI (Dong et al., 2024) and in-plane segmented 3D multi-slab sequences (Z. Li et al., 2024) offer comparable high-fidelity sub-millimeter resolution. Furthermore, the advent of next-generation hardware, such as the Connectome 2.0 (Huang et al., 2021), Next-generation 7T scanners (Feinberg et al., 2023), along with clinically approved systems like the Siemens Cima.X (Kara et al., 2024) is making high-resolution dMRI acquisition increasingly feasible. Second, multi-shell data using a b-value higher than 2000 s/mm^2^ is preferable, since multi-tissue multi-fiber model like MSMT-CSD is capable of separating different tissue and axon compartments and less prone to partial volume effect. When high-resolution acquisition is impractical or dMRI data is retrospectively analyzed, post-processing approaches less sensitive to resolution changes can help yield SAF results closer to those achieved with native high-resolution data. Following approaches given particular acquisition parameters are recommended: for single-shell low b-value data (approximately b = 1000 s/mm²) in DTI, up-sample the dMRI data and use probabilistic tractography with anatomical constraints (Low B, Sing, Prob, w/o SIFT, w/ ACT) appears optimal; for single-shell medium-to-high b-value data (b=2000 s/mm² or higher) and multi-shell data, use CSD-based probabilistic tractography with SIFT and ACT (High B, Xing, Prob, w/ SIFT, w/ ACT and Multi B, Xing, Prob, w/ SIFT, w/ ACT). Finally, researchers should exercise particular caution when interpreting SAF reconstructions originating from cortical regions identified as highly susceptible to resolution effects such as pars opercularis, precuneus and superior temporal (Figure 7, Supplementary Information Figure S5, S6), especially when imaging resolution is limited.

Several limitations highlight avenues for future research. First, the lack of ground truth for SAF reconstruction necessitates validation studies, potentially using tracer injections in post-mortem brains. Second, this study focused on healthy adults. Future work could investigate whether it enhances the sensitivity of SAF and connectivity related biomarkers in patient populations, such as those with neurodegenerative diseases, and in healthy and diseased children. Third, while we evaluated commonly used tractography algorithms for practical relevance, future studies would incorporate emerging surface-based methods (Gahm & Shi, 2019; Y. Li et al., 2024; Nie et al., 2024; Shastin et al., 2022) to provide a more comprehensive understanding of resolution effects and to potentially improve SAF reconstruction accuracy.

## Conclusions

Our study systematically quantified the resolution impact on SAF reconstruction and structural connectivity estimation using multiple-resolution dMRI dataset for 20 healthy subjects. Lower resolution leads to SAF underestimation, especially for simpler models (e.g., DTI model) and basic tractography methods (deterministic, without ACT or SIFT). Data up-sampling prior to tractography partially restores reconstruction accuracy in most cases. High-resolution multi-shell acquisitions for optimal SAF reconstruction is recommended. When unavailable, data up-sampling with DTI-based probabilistic tracking with ACT is advised for single-shell low b-value data, while CSD-based probabilistic tracking with SIFT and ACT is preferable for single-shell high b-value or multi-shell data. This study provides guidelines for improving SAF mapping accuracy, advancing the study of brain connectivity in neuroscience and clinical applications.

## Data and Code Availability

The empirical data are available on request from the corresponding author. The simulation data from 20 subjects are provided by the Human Connectome Project WU-Minn Consortium and are all available via public database at https://www.humanconnectome.org.

The codes for gradient nonlinearity correction are available at https://github.com/ksubramz/gradunwarp. The software used for data processing is also publicly available: FreeSurfer (https://surfer.nmr.mgh.harvard.edu/fswiki/DownloadAndInstall); FMRIB Software Library (FSL) (https://fsl.fmrib.ox.ac.uk/fsl/docs/#/install/index); MRtrix3 (https://www.mrtrix.org/download/); GRETNA (https://www.nitrc.org/frs/?group_id=668). The gSlider sequence and its reconstruction scripts can be accessed through the Collaborating to Commercialize and Publish (C2P) platform (https://www.nmr.mgh.harvard.edu/c2p).

## Author Contributions

Jialan Zheng: Conceptualization, Data curation, Formal analysis, Investigation, Methodology, Software, Validation, Visualization, Writing – original draft; Writing – review & editing

Ziyu Li: Conceptualization, Methodology, Supervision, Validation, Writing – review & editing Wen Zhong: Data curation, Investigation, Methodology

Ziang Wang: Conceptualization

Zihan Li: Conceptualization, Software, Writing – review & editing Hongjia Yang: Conceptualization, Funding acquisition, Resources Mingxuan Liu: Conceptualization

Xiaozhi Cao: Methodology, Resources Congyu Liao: Methodology, Resources

David H. Salat: Conceptualization, Methodology, Supervision Susie Y. Huang: Conceptualization, Methodology, Supervision

Qiyuan Tian: Conceptualization, Funding acquisition, Methodology, Project administration, Resources,

Software, Supervision, Validation, Writing – original draft, Writing – review & editing

## Acknowledgements

We are grateful to Cong Yang for assisting with the scanning procedures and to the 20 subjects who provided their brain imaging data. We also thank the Human Connectome Project (HCP) and its participants for their invaluable time and effort in creating this rich imaging resource.

