## Supplementary Information for "Effects of diffusion MRI spatial resolution on human brain short-range association fiber reconstruction and structural connectivity estimation"

**Table S1. Tracking parameters for empirical data**

| Method | Tracking Algorithm | Seed Number | Seed Location | Step Size (mm) | Angle (°) | Cutoff |
|---|---|---|---|---|---|---|
| Low B, Sing, w/o SIFT, Det, w/o ACT | Tensor_Det | 5M | Whole brain | 0.096 | 60 | 0.1 |
| Low B, Sing, w/o SIFT, Det, w/ ACT | Tensor_Det | 5M | GM-WM interface | 0.096 | 60 | 0.05 |
| Low B, Sing, w/o SIFT, Prob, w/o ACT | Tensor_Prob | 5M | Whole brain | 0.096 | 60 | 0.1 |
| Low B, Sing, w/o SIFT, Prob, w/ ACT | Tensor_Prob | 5M | GM-WM interface | 0.096 | 60 | 0.05 |
| High B, Xing, w/o SIFT, Det, w/o ACT | SD_Stream | 5M | Whole brain | 0.096 | 60 | 0.1 |
| High B, Xing, w/o SIFT, Det, w/ ACT | SD_Stream | 5M | GM-WM interface | 0.096 | 60 | 0.05 |
| High B, Xing, w/o SIFT, Prob, w/o ACT | iFOD2 | 5M | Whole brain | 0.48 | 45 | 0.1 |
| High B, Xing, w/o SIFT, Prob, w/ ACT | iFOD2 | 5M | GM-WM interface | 0.48 | 45 | 0.05 |
| High B, Xing, w/ SIFT, Det, w/o ACT | SD_Stream | 5M | Whole brain | 0.096 | 60 | 0.1 |
| High B, Xing, w/ SIFT, Det, w/ ACT | SD_Stream | 5M | GM-WM interface | 0.096 | 60 | 0.05 |
| High B, Xing, w/ SIFT, Prob, w/o ACT | iFOD2 | 5M | Whole brain | 0.48 | 45 | 0.1 |
| High B, Xing, w/ SIFT, Prob, w/ ACT | iFOD2 | 5M | GM-WM interface | 0.48 | 45 | 0.05 |
| Multi B, Xing, w/o SIFT, Det, w/o ACT | SD_Stream | 5M | Whole brain | 0.096 | 60 | 0.1 |
| Multi B, Xing, w/o SIFT, Det, w/ ACT | SD_Stream | 5M | GM-WM interface | 0.096 | 60 | 0.05 |
| Multi B, Xing, w/o SIFT, Prob, w/o ACT | iFOD2 | 5M | Whole brain | 0.48 | 45 | 0.1 |
| Multi B, Xing, w/o SIFT, Prob, w/ ACT | iFOD2 | 5M | GM-WM interface | 0.48 | 45 | 0.05 |
| Multi B, Xing, w/ SIFT, Det, w/o ACT | SD_Stream | 5M | Whole brain | 0.096 | 60 | 0.05 |
| Multi B, Xing, w/ SIFT, Det, w/ ACT | SD_Stream | 5M | GM-WM interface | 0.096 | 60 | 0.05 |
| Multi B, Xing, w/ SIFT, Prob, w/o ACT | iFOD2 | 5M | Whole brain | 0.48 | 45 | 0.1 |
| Multi B, Xing, w/ SIFT, Prob, w/ ACT | iFOD2 | 5M | GM-WM interface | 0.48 | 45 | 0.05 |

Abbreviations: Low B = single-shell with low b-value; High B = single-shell with high b-value; Multi B = multiple shells with both low and high b-values; Sing = single-fiber model; Xing = crossing-fiber model; Det=deterministic; Prob=probabilistic; SIFT = Spherical-deconvolution Informed Filtering of Tractograms; ACT = anatomical constraints tractography; GM = gray matter; WM = white matter.

**Table S2. Tracking parameters for simulation data**

| Method | Tracking Algorithm | Seed Number | Seed Location | Step Size (mm) | Angle (°) | Cutoff |
|---|---|---|---|---|---|---|
| Low B, Sing, w/o SIFT, Det, w/o ACT | Tensor_Det | 5M | Whole brain | 0.125 | 60 | 0.1 |
| Low B, Sing, w/o SIFT, Det, w/ ACT | Tensor_Det | 5M | GM-WM interface | 0.125 | 60 | 0.05 |
| Low B, Sing, w/o SIFT, Prob, w/o ACT | Tensor_Prob | 5M | Whole brain | 0.125 | 60 | 0.1 |
| Low B, Sing, w/o SIFT, Prob, w/ ACT | Tensor_Prob | 5M | GM-WM interface | 0.125 | 60 | 0.05 |
| High B, Xing, w/o SIFT, Det, w/o ACT | SD_Stream | 5M | Whole brain | 0.125 | 60 | 0.1 |
| High B, Xing, w/o SIFT, Det, w/ ACT | SD_Stream | 5M | GM-WM interface | 0.125 | 60 | 0.05 |
| High B, Xing, w/o SIFT, Prob, w/o ACT | iFOD2 | 5M | Whole brain | 0.625 | 45 | 0.1 |
| High B, Xing, w/o SIFT, Prob, w/ ACT | iFOD2 | 5M | GM-WM interface | 0.625 | 45 | 0.05 |
| High B, Xing, w/ SIFT, Det, w/o ACT | SD_Stream | 5M | Whole brain | 0.125 | 60 | 0.1 |
| High B, Xing, w/ SIFT, Det, w/ ACT | SD_Stream | 5M | GM-WM interface | 0.125 | 60 | 0.05 |
| High B, Xing, w/ SIFT, Prob, w/o ACT | iFOD2 | 5M | Whole brain | 0.625 | 45 | 0.1 |
| High B, Xing, w/ SIFT, Prob, w/ ACT | iFOD2 | 5M | GM-WM interface | 0.625 | 45 | 0.05 |
| Multi B, Xing, w/o SIFT, Det, w/o ACT | SD_Stream | 5M | Whole brain | 0.125 | 60 | 0.1 |
| Multi B, Xing, w/o SIFT, Det, w/ ACT | SD_Stream | 5M | GM-WM interface | 0.125 | 60 | 0.05 |
| Multi B, Xing, w/o SIFT, Prob, w/o ACT | iFOD2 | 5M | Whole brain | 0.625 | 45 | 0.1 |
| Multi B, Xing, w/o SIFT, Prob, w/ ACT | iFOD2 | 5M | GM-WM interface | 0.625 | 45 | 0.05 |
| Multi B, Xing, w/ SIFT, Det, w/o ACT | SD_Stream | 5M | Whole brain | 0.125 | 60 | 0.05 |
| Multi B, Xing, w/ SIFT, Det, w/ ACT | SD_Stream | 5M | GM-WM interface | 0.125 | 60 | 0.05 |
| Multi B, Xing, w/ SIFT, Prob, w/o ACT | iFOD2 | 5M | Whole brain | 0.625 | 45 | 0.1 |
| Multi B, Xing, w/ SIFT, Prob, w/ ACT | iFOD2 | 5M | GM-WM interface | 0.625 | 45 | 0.05 |

Abbreviations: Low B = single-shell with low b-value; High B = single-shell with high b-value; Multi B = multiple shells with both low and high b-values; Sing = single-fiber model; Xing = crossing-fiber model; Det=deterministic; Prob=probabilistic; SIFT = Spherical-deconvolution Informed Filtering of Tractograms; ACT = anatomical constraints tractography; GM = gray matter; WM = white matter.

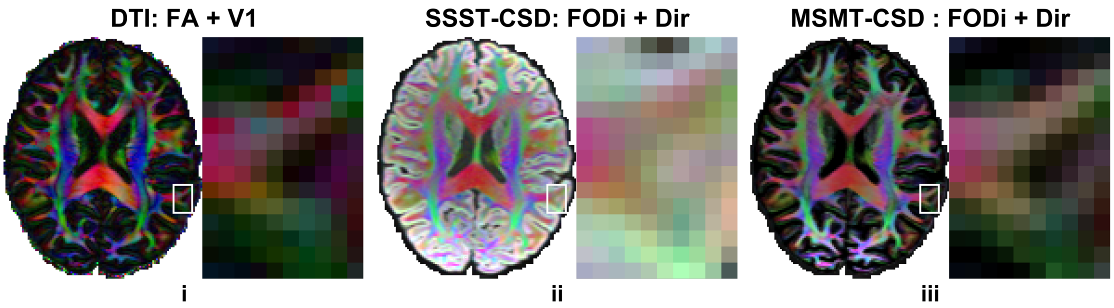


**Figure S1. HCP data quality.** Axial slices from a representative subject with fitted results from three diffusion models (i: DTI, ii: SSST-CSD, iii: MSMT-CSD) at 1.25 mm iso. resolution are displayed. The fractional anisotropy (FA) map color-coded by primary vector (V1) (i) and the fiber orientation distribution integral (FODi) map color-coded by the overall fiber direction (ii, iii) are shown for DTI and CSD methods (red: left–right; green: anterior–posterior; blue: superior–inferior), respectively.

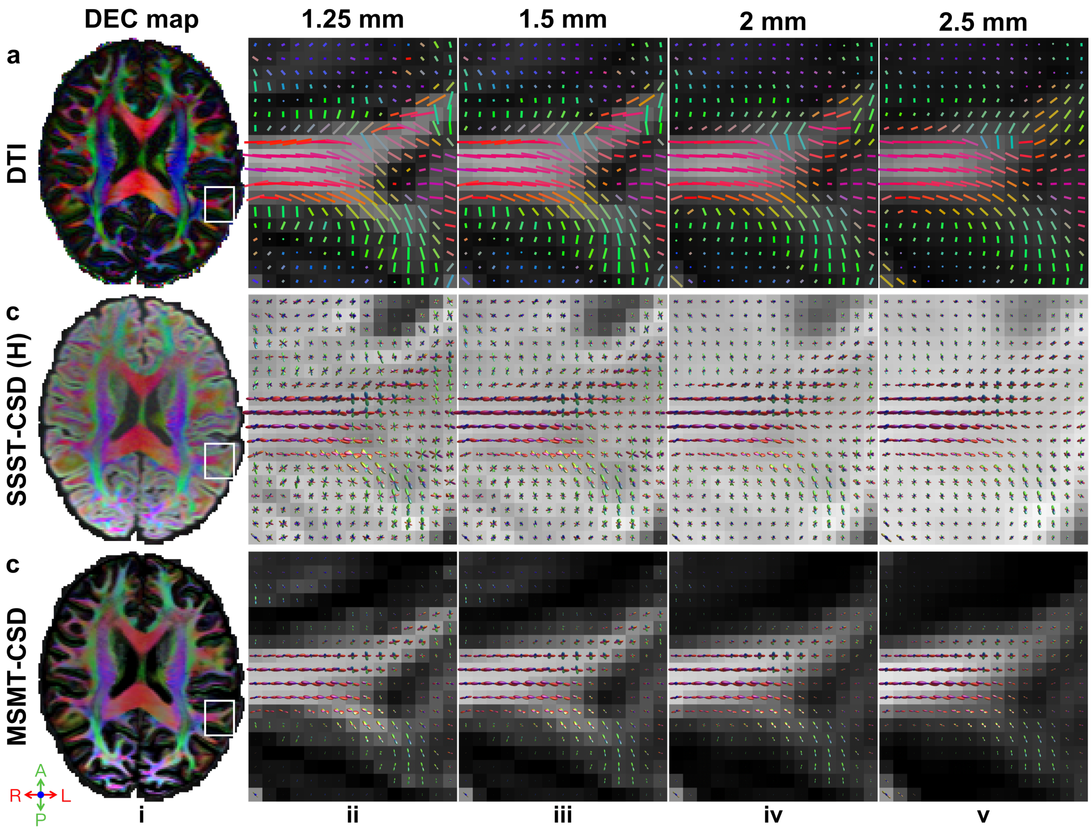


**Figure S2. Axon orientations from simulation data.** Primary fiber direction encoded color (DEC) maps (red: left–right; green: anterior–posterior; blue: superior–inferior) at 1.25 mm iso. resolution from a representative subject (i) are displayed for three methods including DTI (a, i), SSST-CSD (b, i), and MSMT-CSD (c, i), with a region of interest (white boxes) containing a gyrus, subcortical white matter, and their interface shown in enlarged views overlaid on FA (a), and FOD integral (b, c) maps across four different spatial resolutions (ii: 1.25 mm, iii: 1.5 mm, iv: 2 mm, v: 2.5 mm).

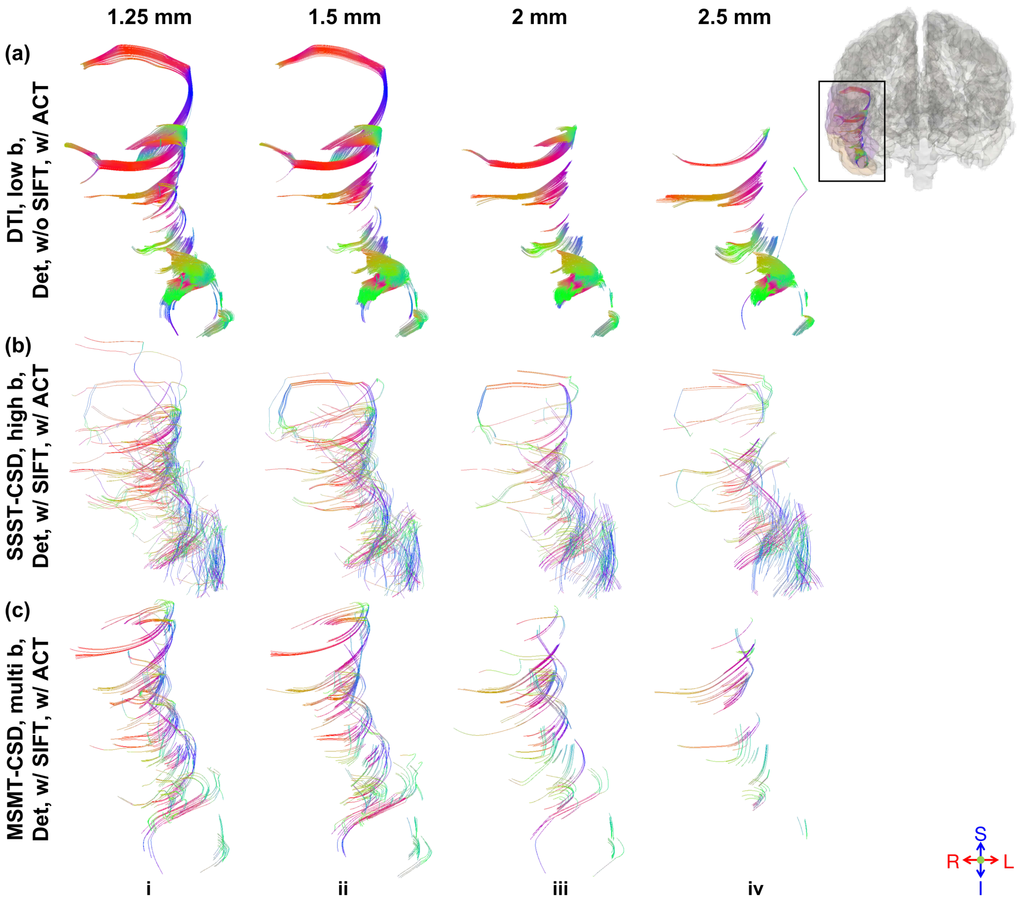


**Figure S3. Simulation data tractography results.** U-shaped SAFs connecting the right middle and superior temporal gyri (black box) reconstructed using three modeling methods (a: DTI on low b-value single-shell data, b: SSST-CSD on high b-value single-shell data with SIFT, c: MSMT-CSD on multi b-value data with SIFT) and deterministic tractography with ACT across four different spatial resolutions (i: 1.25 mm, ii: 1.5 mm, iii: 2 mm, iv: 2.5 mm) are displayed.

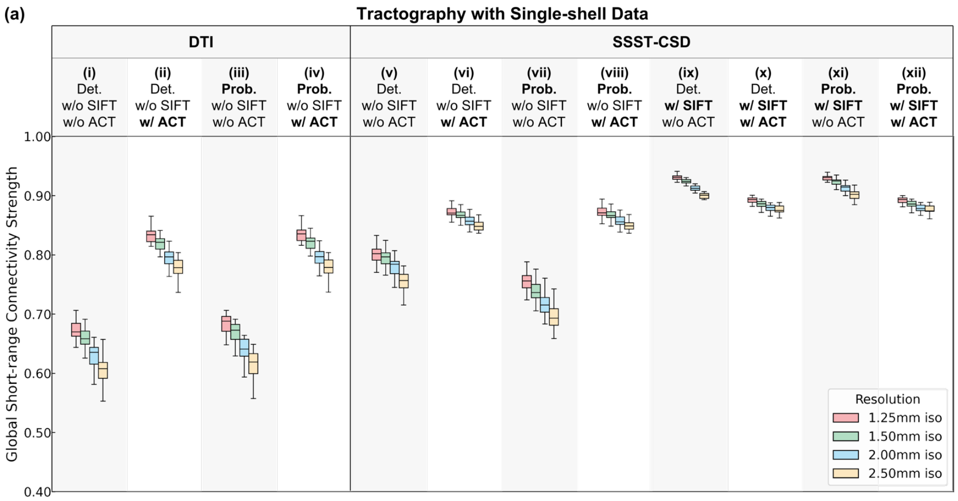


| | (i) | (ii) | (iii) | (iv) | (v) | (vi) | (vii) | (viii) | (ix) | (x) | (xi) | (xii) |
|---|---|---|---|---|---|---|---|---|---|---|---|---|
| | -2.04%*** | -1.52%*** | -2.32%*** | -1.57%*** | -0.64%*** | -0.65%*** | -2.05%*** | -0.62%*** | -0.67%*** | -0.79%*** | -0.67%*** | -0.74%*** |
| | -6.16%*** | -4.31%*** | -6.50%*** | -4.45%*** | -2.64%*** | -1.87%*** | -4.74%*** | -1.87%*** | -1.90%*** | -1.48%*** | -1.81%*** | -1.48%*** |
| | -10.00%*** | -6.53%*** | -10.17%*** | -6.67%*** | -5.55%*** | -2.72%*** | -7.63%*** | -2.66%*** | -3.23%*** | -1.74%*** | -3.07%*** | -1.68%*** |
| | -4.20%*** | -2.83%*** | -4.27%*** | -2.92%*** | -2.02%*** | -1.22%*** | -2.75%*** | -1.25%*** | -1.24%*** | -0.69%*** | -1.14%*** | -0.74%*** |
| | -8.13%*** | -5.09%*** | -8.03%*** | -5.18%*** | -4.94%*** | -2.08%*** | -5.70%*** | -2.05%*** | -2.58%*** | -0.95%*** | -2.41%*** | -0.95%*** |
| | -4.10%*** | -2.32%*** | -3.93%*** | -2.33%*** | -2.98%*** | -0.87%*** | -3.04%*** | -0.81%*** | -1.35%*** | -0.26%* | -1.28%*** | -0.20%* |

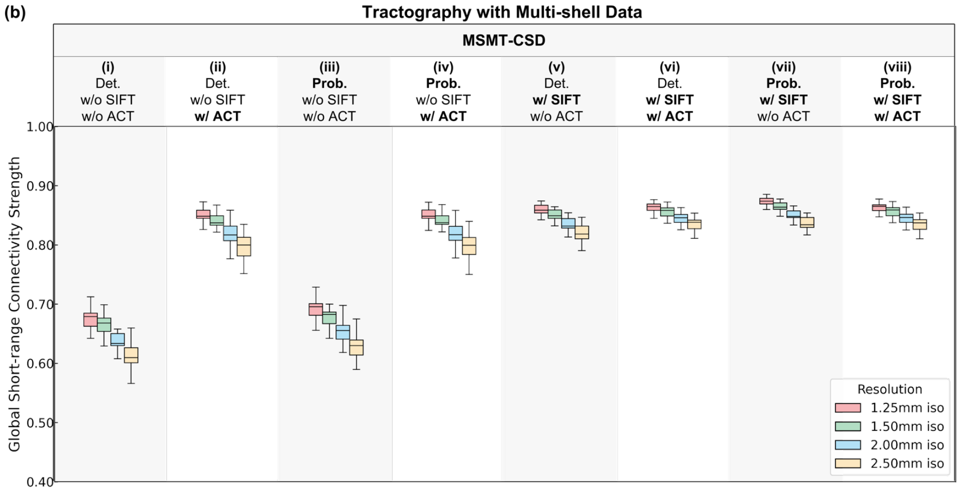


| | (i) | (ii) | (iii) | (iv) | (v) | (vi) | (vii) | (viii) |
|---|---|---|---|---|---|---|---|---|
| | -1.60%*** | -1.24%*** | -1.88%*** | -1.17%*** | -1.02%*** | -0.73%*** | -1.03%*** | -0.67%*** |
| | -5.47%*** | -4.07%*** | -5.54%*** | -4.01%*** | -2.88%*** | -2.18%*** | -2.73%*** | -2.18%*** |
| | -9.25%*** | -6.34%*** | -9.24%*** | -6.28%*** | -4.66%*** | -3.38%*** | -4.29%*** | -3.38%*** |
| | -3.94%*** | -2.86%*** | -3.72%*** | -2.88%*** | -1.88%*** | -1.46%*** | -1.71%*** | -1.52%*** |
| | -7.78%*** | -5.16%*** | -7.50%*** | -5.18%*** | -3.67%*** | -2.66%*** | -3.29%*** | -2.72%*** |
| | -4.00%*** | -2.37%*** | -3.92%*** | -2.37%*** | -1.83%*** | -1.22%*** | -1.60%*** | -1.22%*** |

**Figure S4. Simulation data GSCS.** Box plots for GSCS from different tractography methods using single-shell (a) and multi-shell (b) data at three native spatial resolutions (red: 0.96 mm iso., green: 1.5 mm iso., blue: 2 mm iso.) and two nominally high 0.96 mm iso. resolution up-sampled from 1.5 mm and 2 mm iso. resolution (hatched green: 1.5 mm up-sampled, hatched blue: 2 mm up-sampled) display the distribution (e.g., median, interquartile range, and range) of GSCS from 20 subjects in the upper panel. The lower table lists the relative difference of GSCS at lower resolution compared to the GSCS at higher resolution (each row) for different tractography methods (each column), with asterisks denoting significance levels (*: $p < 0.05$, **: $p < 0.01$, ***: $p < 0.001$). The color of each table cell indicates the magnitude and direction of the GSCS difference (blue: decrease, red: increase)

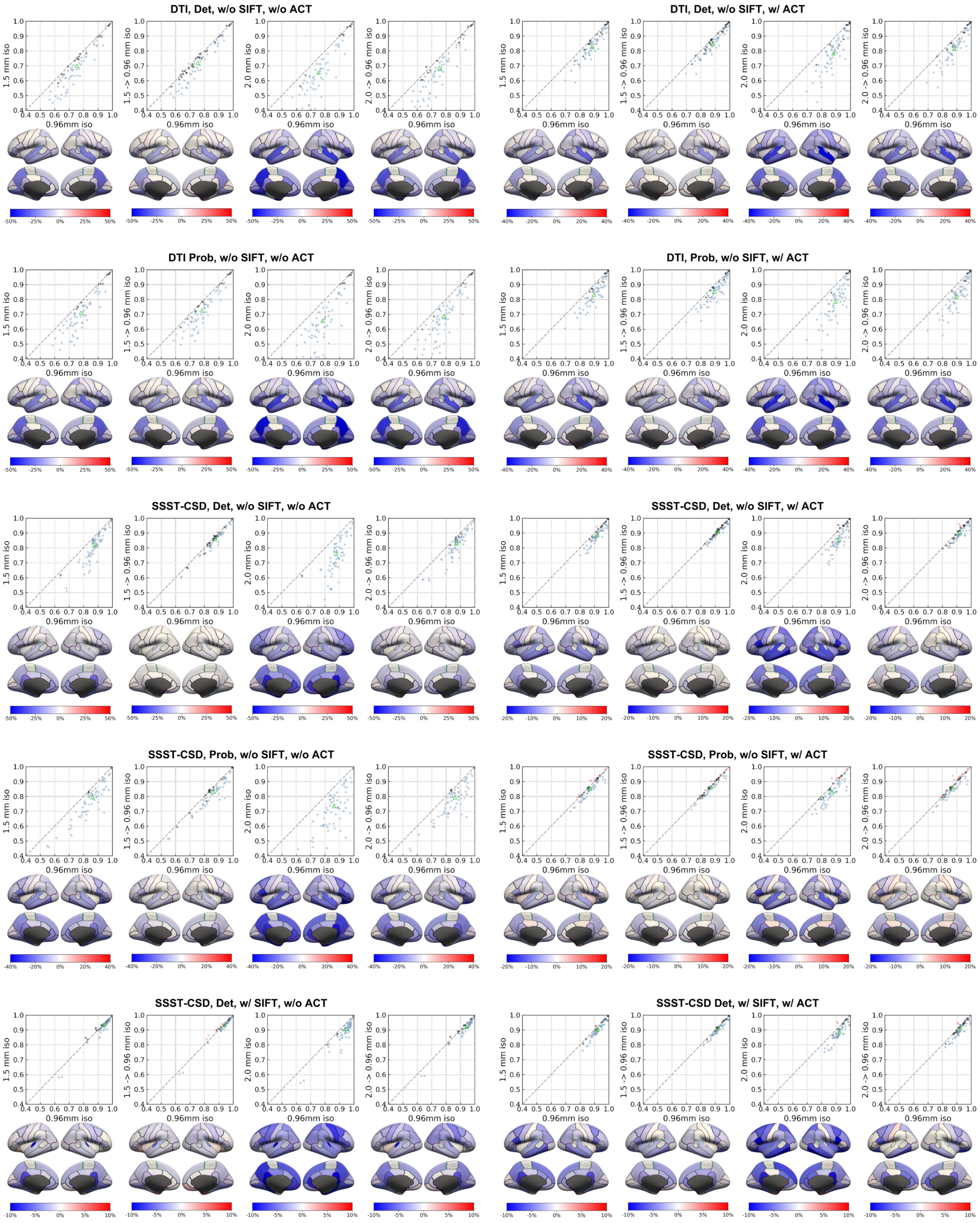
DTI, Det, w/o SIFT, w/o ACT
DTI, Det, w/o SIFT, w/ ACT
DTI Prob, w/o SIFT, w/o ACT
DTI, Prob, w/o SIFT, w/ ACT
SSST-CSD, Det, w/o SIFT, w/o ACT
SSST-CSD, Det, w/o SIFT, w/ ACT
SSST-CSD, Prob, w/o SIFT, w/o ACT
SSST-CSD, Prob, w/o SIFT, w/ ACT
SSST-CSD, Det, w/ SIFT, w/o ACT
SSST-CSD Det, w/ SIFT, w/ ACT
1.5 mm iso
1.5 -> 0.96 mm iso
2.0 mm iso
2.0 -> 0.96 mm iso
0.96mm iso

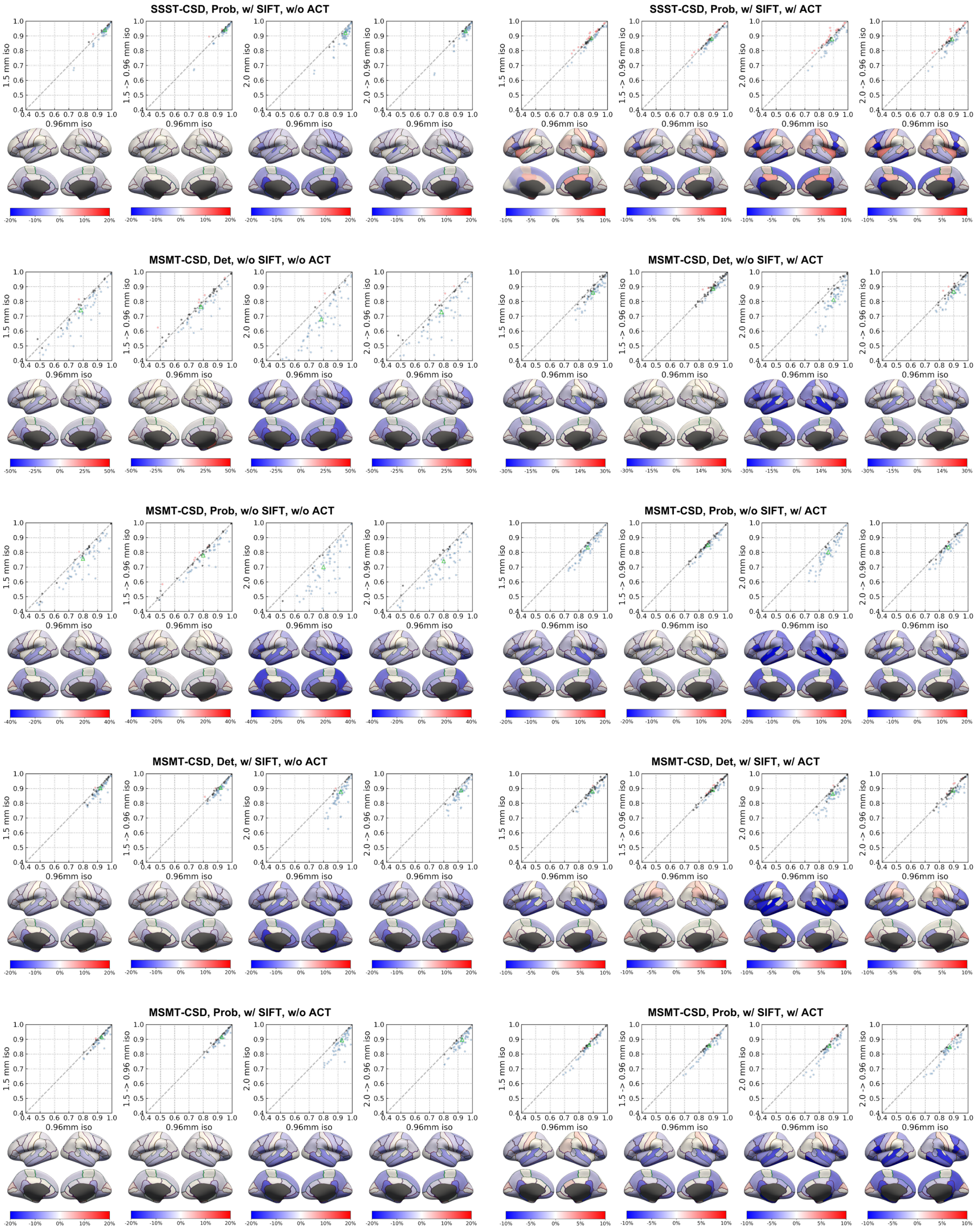
SSST-CSD, Prob, w/ SIFT, w/o ACT
SSST-CSD, Prob, w/ SIFT, w/ ACT
MSMT-CSD, Det, w/o SIFT, w/o ACT
MSMT-CSD, Det, w/o SIFT, w/ ACT
MSMT-CSD, Prob, w/o SIFT, w/o ACT
MSMT-CSD, Prob, w/o SIFT, w/ ACT
MSMT-CSD, Det, w/ SIFT, w/o ACT
MSMT-CSD, Det, w/ SIFT, w/ ACT
MSMT-CSD, Prob, w/ SIFT, w/o ACT
MSMT-CSD, Prob, w/ SIFT, w/ ACT
1.5 mm iso
1.5 -> 0.96 mm iso
2.0 mm iso
2.0 -> 0.96 mm iso
0.96mm iso

**Figure S5. Comprehensive empirical data RSCS.** Scatter plots comparing RSCS at high resolution (0.96mm, x-axis) versus at various low-resolution conditions (1.5 mm native, 1.5 mm up-sampled to 0.96 mm, 2 mm native, and 2 mm up-sampled to 0.96 mm; y-axis) are displayed with each point representing a cortical region, color representing the difference and its significance (blue = decrease, red = increase, black = none), and the green triangle representing GSCS for various tractography methods (detailed by panel titles). The difference of RSCS at each low-resolution condition compared to RSCS at high resolution for each cortical region is displayed on inflated surfaces below the corresponding scatter plot, with color and its intensity representing the pattern.

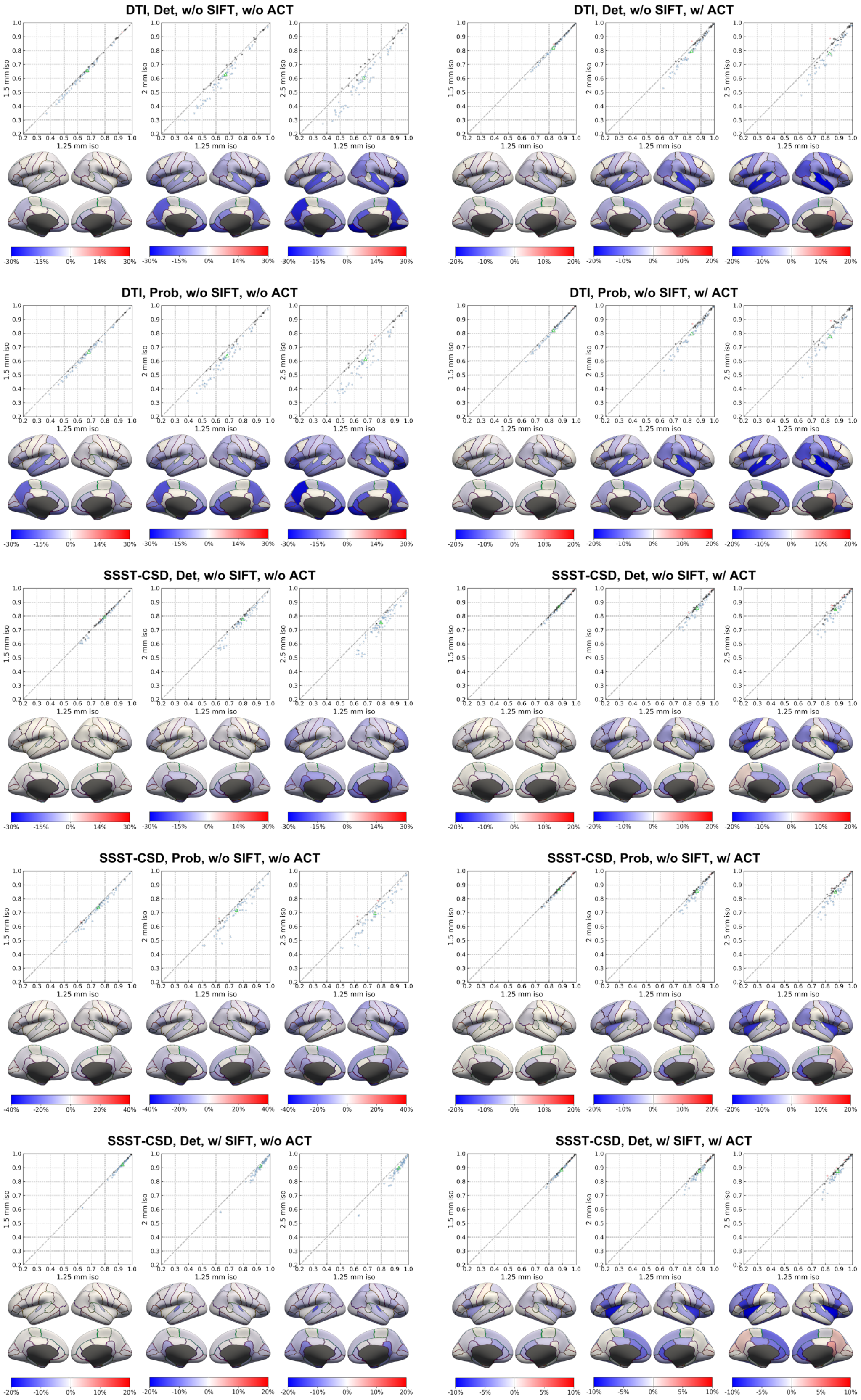
DTI, Det, w/o SIFT, w/o ACT
DTI, Det, w/o SIFT, w/ ACT
DTI, Prob, w/o SIFT, w/o ACT
DTI, Prob, w/o SIFT, w/ ACT
SSST-CSD, Det, w/o SIFT, w/o ACT
SSST-CSD, Det, w/o SIFT, w/ ACT
SSST-CSD, Prob, w/o SIFT, w/o ACT
SSST-CSD, Prob, w/o SIFT, w/ ACT
SSST-CSD, Det, w/ SIFT, w/o ACT
SSST-CSD, Det, w/ SIFT, w/ ACT
1.5 mm iso
2 mm iso
2.5 mm iso
1.25 mm iso

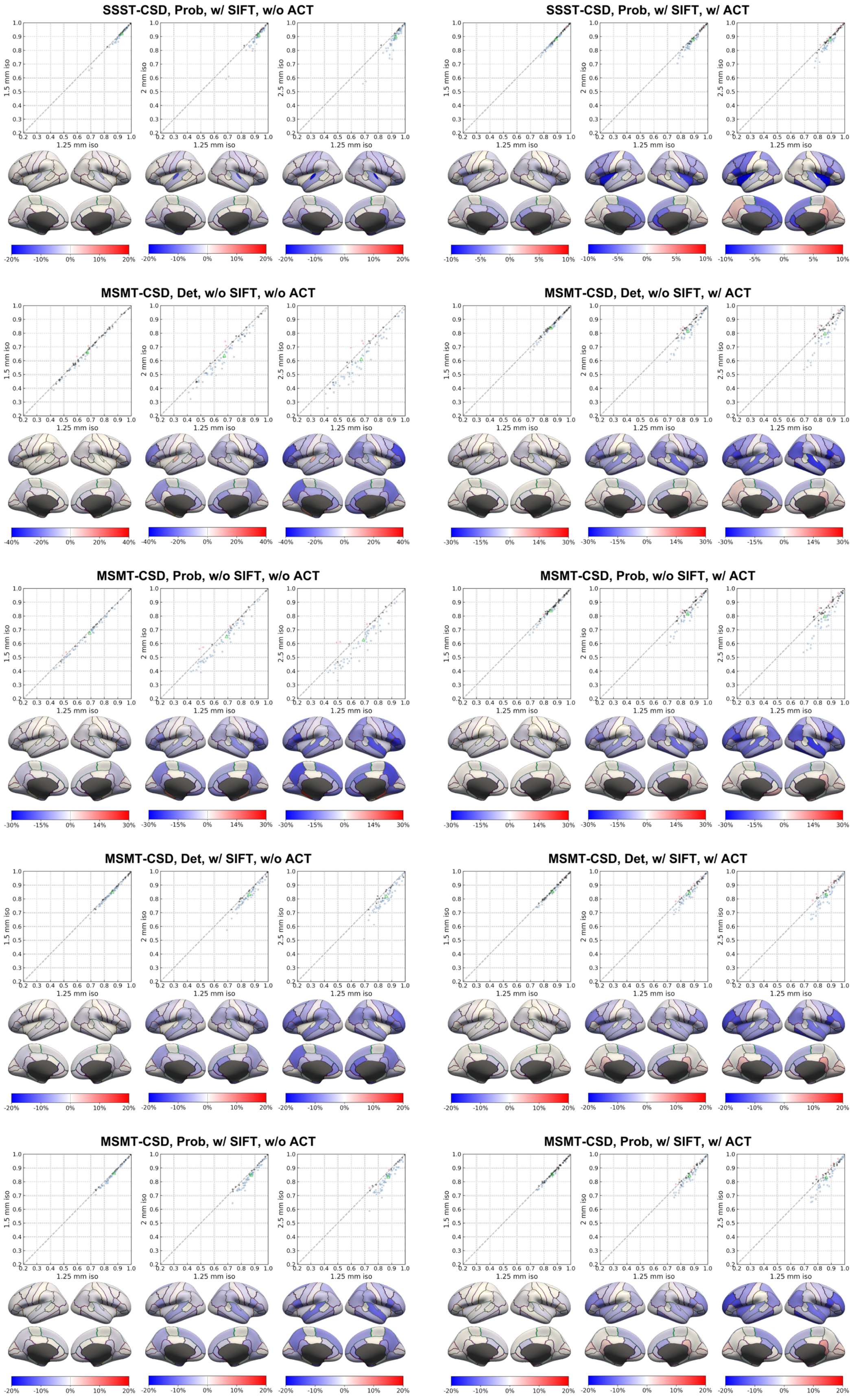
SSST-CSD, Prob, w/ SIFT, w/o ACT
SSST-CSD, Prob, w/ SIFT, w/ ACT
MSMT-CSD, Det, w/o SIFT, w/o ACT
MSMT-CSD, Det, w/o SIFT, w/ ACT
MSMT-CSD, Prob, w/o SIFT, w/o ACT
MSMT-CSD, Prob, w/o SIFT, w/ ACT
MSMT-CSD, Det, w/ SIFT, w/o ACT
MSMT-CSD, Det, w/ SIFT, w/ ACT
MSMT-CSD, Prob, w/ SIFT, w/o ACT
MSMT-CSD, Prob, w/ SIFT, w/ ACT
1.25 mm iso
1.5 mm iso
2 mm iso
2.5 mm iso
-20% -10% 0% 10% 20%
-10% -5% 0% 5% 10%
-40% -20% 0% 20% 40%
-30% -15% 0% 14% 30%

**Figure S6. Comprehensive simulation data RSCS.** Scatter plots comparing RSCS at high resolution (1.25 mm, x-axis) versus at various low-resolution conditions (1.5 mm, 2 mm, and 2.5 mm; y-axis) are displayed with each point representing a cortical region, color representing the difference and its significance (blue = decrease, red = increase, black = none), and the green triangle representing GSCS for various tractography methods (detailed by panel titles). The difference of RSCS at each low-resolution condition compared to RSCS at high resolution for each cortical region is displayed on inflated surfaces below the corresponding scatter plot, with color and its intensity representing the pattern.

**Small-Worldness Calculation and Results**

Small-Worldness (SW) quantifies the balance between local segregation and global integration of the structural connectome. A small-world brain network exhibits a high clustering coefficient (i.e. brain regions structurally connected to a particular region also tend to be interconnected with each other) and a short characteristic path length (i.e. strong neural pathways can generally be found between any two brain regions). SW is calculated as the ratio of the brain network's small-world properties to those of a random network. SW > 1 indicates that the brain network demonstrates stronger small-world properties compared to a random network. SW metrics were calculated using the GRETNA toolbox in MATLAB (Wang et al., 2015) described as follows.

The structural connectivity matrix $\mathrm{W}$ can be represented as a weighted, undirected graph $\mathrm{G} = (\mathrm{N}, \mathrm{E})$, where $\mathrm{N}$ is the set of nodes (cortical regions as defined by the Desikan-Killiany atlas) and $\mathrm{E}$ is the set of edges. An edge $(\mathrm{i}, \mathrm{j})$ exists between nodes $\mathrm{i}$ and $\mathrm{j}$ if their connectivity strength $\omega_{\mathrm{i,j}} > 0$. SW is calculated as the ratio of the normalized clustering coefficient ($\gamma$) to the normalized characteristic path length ($\lambda$) (Watts & Strogatz, 1998), separately for each hemispheric sub-graph $\mathrm{G_h}$ (where $\mathrm{h} \in \{\mathrm{left}, \mathrm{right}\}$):

$$\mathrm{SW}_h = \frac{\gamma_h}{\lambda_h}. \tag{1}$$

The normalized clustering coefficient $\gamma_\mathrm{h} = \mathrm{C_h}/\mathrm{C_{h,rand}}$ compares the average clustering coefficient $\mathrm{C_h}$ of the hemispheric network $\mathrm{G_h}$ to the average value $\mathrm{C_{h,rand}}$ obtained from an ensemble of random networks. These random networks are generated by rewiring the original graph's edges while preserving the degree sequence (i.e., each node $\mathrm{k} \in \mathrm{N_h}$ maintains its original degree $\mathrm{d_k} = |\{\mathrm{i} \in \mathrm{N_h} \mid \omega_{\mathrm{ki}} > 0\}|$) (Maslov & Sneppen, 2002). The average clustering coefficient $\mathrm{C_h}$ is the mean of the local clustering coefficients $\mathrm{C_k}$ across all nodes $\mathrm{k} \in \mathrm{N_h}$. The local clustering coefficient $\mathrm{C_k}$ for a node $\mathrm{k}$ is calculated as (Onnela et al., 2005):

$$C_k = \frac{2}{d_k(d_k-1)} \sum_{i,j \subset N_k} \left(\widehat{\omega}_{ki}\widehat{\omega}_{ij}\widehat{\omega}_{jk}\right)^{1/3} \tag{2}$$

where $N_k = \{i \in N_h \mid \omega_{ki} > 0\}$ is the set of neighbors of node $k$ within the hemisphere $h$. The term $\widehat{\omega_{ij}} = \omega_{ij}/\max(\omega_h)$ represents edge weights normalized by the maximum weight in the hemispheric network $G_h$.

The normalized characteristic path length $\lambda_h = L_h/L_{h,\text{rand}}$ compares the characteristic path length $L_h$ of the network $G_h$ to the average value $L_{h,\text{rand}}$ obtained from the same ensemble of degree-matched random networks. The characteristic path length $L_h$ is the average shortest path length $l_{ij}$ between all distinct node pairs $i, j \in N_h$. The shortest path length $l_{ij}$ is defined as the minimum sum of inverse edge weights along any path $P_{ij} = (i = v_0, v_1, \ldots, v_k = j)$ connecting nodes $i$ and $j$:

$$l_{ij} = \min_{P_{ij}} \sum_{m=0}^{k-1} \frac{1}{\omega_{v_m v_{m+1}}} \tag{3}$$

where $\omega_{v_m v_{m+1}}$ is the weight of the edge between consecutive nodes $v_m$ and $v_{m+1}$ in the path $P_{ij}$ (Dijkstra, 1959).

To mitigate the influence of potentially spurious weak connections (often false positives), a range of connection sparsity thresholds $p$ (5% to 50% in 5% increments) was applied to each $G_h$ to create $G_{h,p} = (N_h, E_{h,p})$, where $E_{h,p} \subseteq E_h$ such that $|E_{h,p}| = \lceil p \times |E_h| \rceil$ and for any $(u, v) \in E_{h,p}$ and $(x, y) \in E_h \setminus E_{h,p}$, $\omega_{uv} \geq \omega_{xy}$. For each $G_{h,p}$, $SW_{h,p}$ was calculated. This procedure generated a curve of $SW_{h,p}$ as a function of $p$. The area under this $SW_{h,p}$ versus $p$ curve, integrated across the sparsity thresholds from 5% to 50%, was calculated. This integrated value was subsequently divided by 0.45 (corresponding to the span of a 5% to 50% sparsity range) to derive $SW_h$, a summary measure robust to threshold selection. The final SW value for the whole brain graph $G$ was calculated as $(SW_{\text{left}} + SW_{\text{right}})/2$.

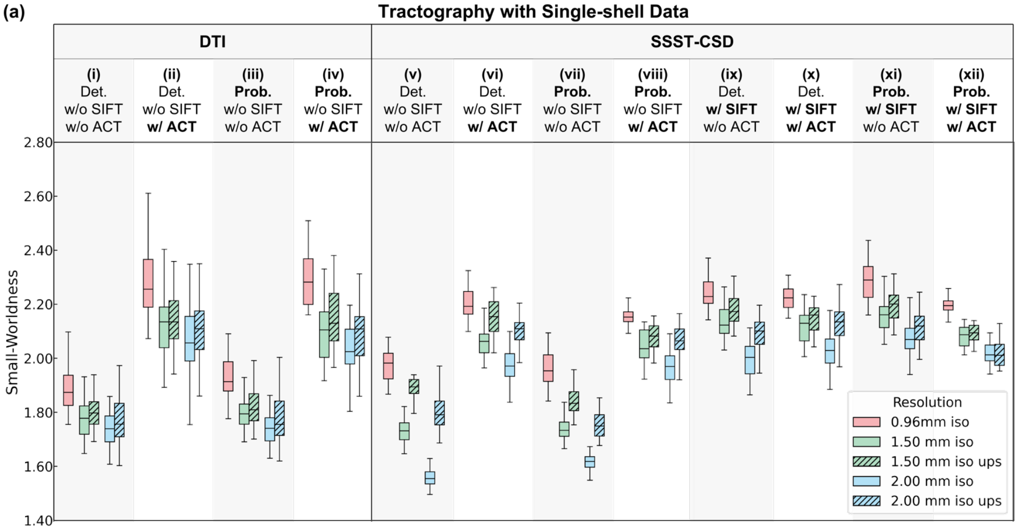


| | (i) | (ii) | (iii) | (iv) | (v) | (vi) | (vii) | (viii) | (ix) | (x) | (xi) | (xii) |
|---|---|---|---|---|---|---|---|---|---|---|---|---|
| | -5.54%*** | -7.17%*** | -6.60%*** | -8.48%*** | -12.04%*** | -6.73%*** | -11.09%*** | -5.18%*** | -5.06%*** | -4.73%*** | -5.48%*** | -5.26%*** |
| | -7.47%*** | -10.08%*** | -9.41%*** | -11.17%*** | -20.65%*** | -11.00%*** | -17.01%*** | -8.65%*** | -10.95%*** | -8.80%*** | -9.60%*** | -8.30%*** |
| | -2.05%** | -3.14%*** | -3.01%*** | -2.93%*** | -9.80%*** | -4.59%*** | -6.65%*** | -3.66%*** | -6.21%*** | -4.27%*** | -4.36%*** | -3.21%*** |
| | -4.08%*** | -5.90%*** | -5.74%*** | -5.99%*** | -3.84%*** | -2.32%** | -6.04%*** | -3.38%*** | -2.78%*** | -3.45%*** | -4.37%*** | -4.86%*** |
| | -6.08%*** | -8.49%*** | -8.19%*** | -9.04%*** | -8.63%*** | -4.53%*** | -10.47%*** | -4.30%*** | -6.71%*** | -4.22%*** | -7.61%*** | -8.25%*** |
| | -2.08%** | -2.76%*** | -2.60%** | -3.25%*** | -4.99%*** | -2.27%*** | -4.71%*** | -0.95% | -4.04%*** | -0.80% | -3.39%*** | -3.56%*** |
| | 1.54%*** | 1.37%** | 0.92%** | 2.73%*** | 9.32%*** | 4.73%*** | 5.69%*** | 1.90%*** | 2.39%*** | 1.35%* | 1.17%** | 0.43% |
| | 1.51%** | 1.77%** | 1.35%* | 2.39%*** | 15.15%*** | 7.27%*** | 7.88%*** | 4.77%*** | 4.76%*** | 5.02%*** | 2.20%*** | 0.06% |

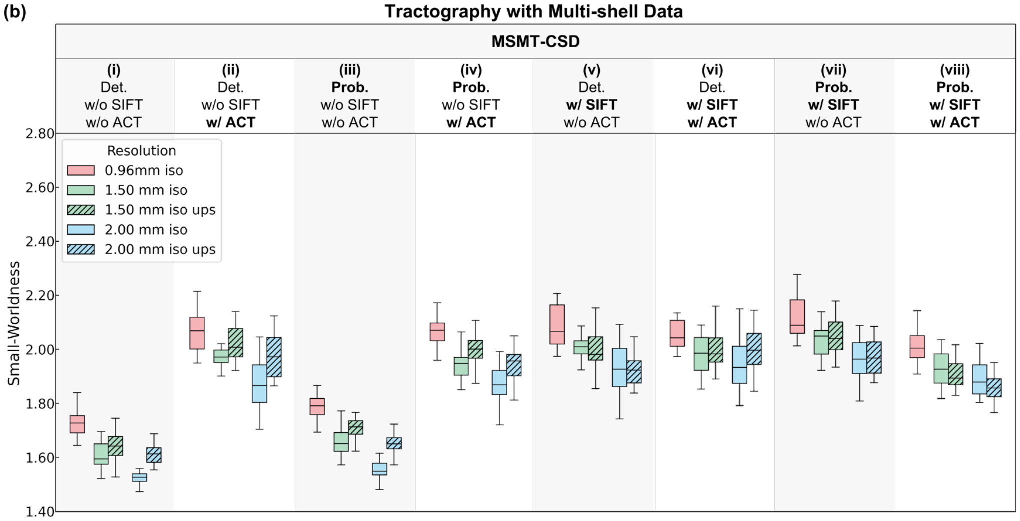


| | (i) | (ii) | (iii) | (iv) | (v) | (vi) | (vii) | (viii) |
|---|---|---|---|---|---|---|---|---|
| | -7.32%*** | -4.71%*** | -7.56%*** | -5.90%*** | -3.67%*** | -3.40%*** | -3.89%*** | -4.19%*** |
| | -11.50%*** | -9.37%*** | -13.42%*** | -9.30%*** | -7.46%*** | -4.97%*** | -7.43%*** | -5.89%*** |
| | -4.50%*** | -4.89%*** | -6.35%*** | -3.62%*** | -3.93%*** | -1.63%* | -3.68%*** | -1.78%*** |
| | -5.36%*** | -2.30%** | -4.42%*** | -3.48%*** | -3.80%*** | -2.50%*** | -3.44%*** | -5.04%*** |
| | -6.88%*** | -4.45%*** | -7.94%*** | -5.91%*** | -7.31%*** | -2.40%** | -6.97%*** | -7.38%*** |
| | -1.61%** | -2.20%** | -3.69%*** | -2.52%*** | -3.65%*** | 0.09% | -3.65%*** | -2.46%*** |
| | 2.12%*** | 2.53%*** | 3.39%*** | 2.57%*** | -0.13% | 0.93%* | 0.47% | -0.89%* |
| | 5.21%*** | 5.43%*** | 6.33%*** | 3.74%*** | 0.16% | 2.70%*** | 0.49% | -1.58%*** |

**Figure S7 Empirical data SW.** Box plots for SW from different tractography methods using single-shell (a) and multi-shell (b) data at three native spatial resolutions (red: 0.96 mm iso., green: 1.5 mm iso., blue: 2 mm iso.) and two nominally high 0.96 mm iso. resolution up-sampled from 1.5 mm and 2 mm iso. resolution (hatched green: 1.5 mm up-sampled, hatched blue: 2 mm up-sampled) display the distribution (i.e., median, interquartile range, and range) of SW from 20 subjects in the upper panel. The lower table lists the relative difference of SW at lower resolution compared to the SW at higher resolution (each row) for different tractography methods (each column), with asterisks denoting significance levels (*: $p < 0.05$, **: $p < 0.01$, ***: $p < 0.001$). The color of each table cell indicates the magnitude and direction of the GSCS difference (blue: decrease, red: increase)

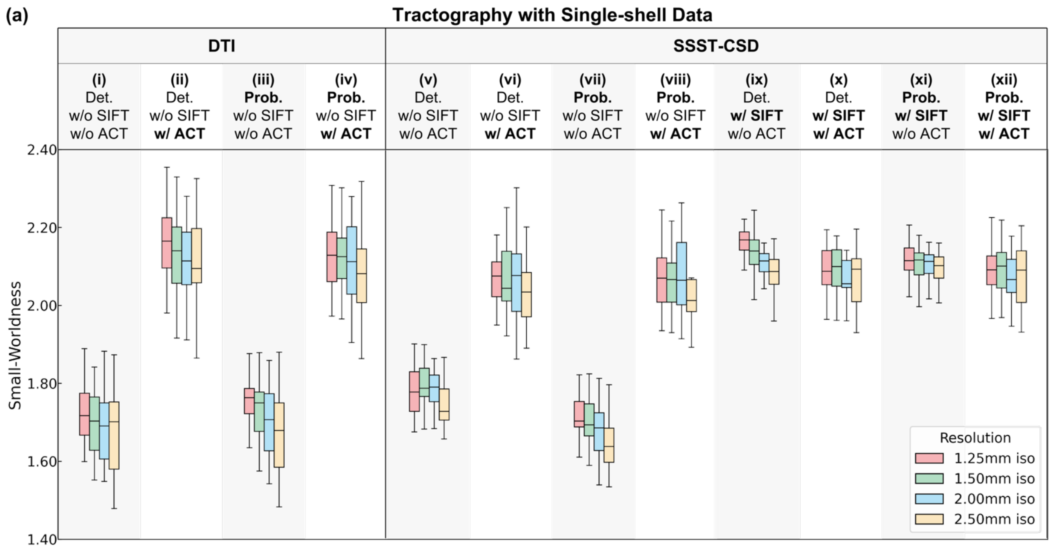


| (i) | (ii) | (iii) | (iv) | (v) | (vi) | (vii) | (viii) | (ix) | (x) | (xi) | (xii) |
|---|---|---|---|---|---|---|---|---|---|---|---|
| -1.59%*** | -1.19%** | -0.86%* | -0.25% | 0.86%** | -0.08% | -1.40%** | -0.19% | -1.07%** | 0.01% | -0.61% | -0.01% |
| -2.39%** | -1.96%*** | -2.81%*** | -1.18%* | 0.49% | -0.52% | -2.65%*** | 0.19% | -2.44%*** | -1.36%** | -0.86% | -0.72% |
| -3.33%** | -2.31%** | -4.06%*** | -2.34%** | -2.00%** | -1.40% | -4.70%*** | -1.54% | -3.53%*** | -0.85% | -1.35%* | -0.88% |
| -0.81% | -0.78% | -1.97%*** | -0.93% | -0.37% | -0.44% | -1.26%** | 0.38% | -1.39%** | -1.38%** | -0.25% | -0.71%* |
| -1.76% | -1.13% | -3.23%*** | -2.10%** | -2.84%*** | -1.33% | -3.35%*** | -1.35% | -2.48%*** | -0.87% | -0.74%* | -0.86% |
| -0.96% | -0.36% | -1.28%** | -1.17%* | -2.48%*** | -0.89% | -2.11%*** | -1.73%* | -1.11%** | 0.52% | -0.49% | -0.16% |

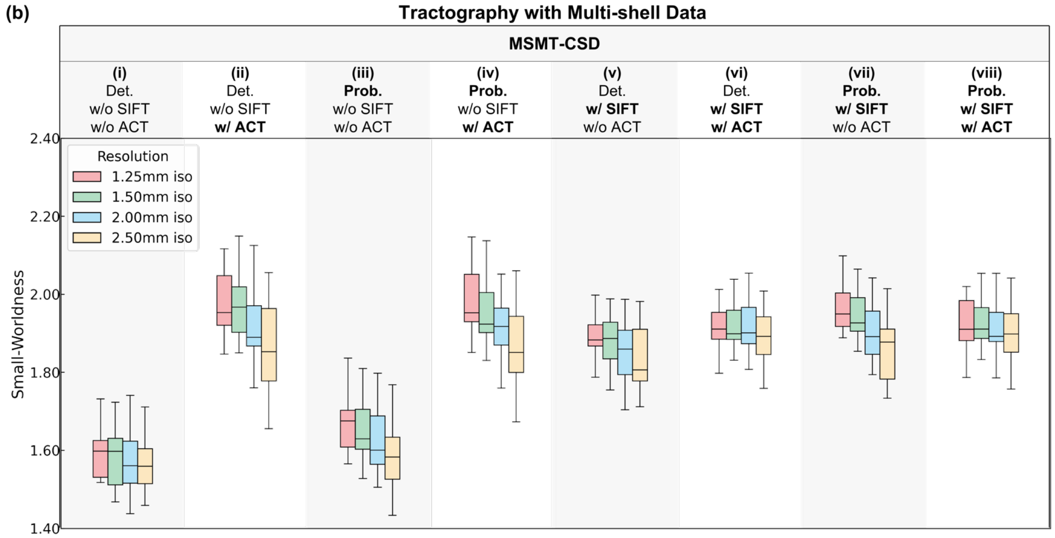


| (i) | (ii) | (iii) | (iv) | (v) | (vi) | (vii) | (viii) |
|---|---|---|---|---|---|---|---|
| -0.75%* | -0.57% | -1.13%** | -0.60% | -0.76%** | 0.52% | -1.01%* | 0.21% |
| -1.66%** | -3.06%*** | -2.72%*** | -2.96%*** | -1.88%** | 0.10% | -3.12%*** | -0.39% |
| -1.98%* | -5.56%*** | -5.13%*** | -5.58%*** | -3.20%*** | -1.22% | -5.30%*** | -1.35% |
| -0.92% | -2.51%*** | -1.61%** | -2.37%** | -1.12% | -0.42% | -2.13%*** | -0.59% |
| -1.24% | -5.02%*** | -4.05%*** | -5.01%*** | -2.45%** | -1.73%** | -4.33%*** | -1.56%* |
| -0.33% | -2.58%** | -2.47%*** | -2.70%*** | -1.34%** | -1.32%** | -2.25%*** | -0.97% |

**Figure S8. Simulation data SW.** Box plots for SW from different tractography methods using single-shell (a) and multi-shell (b) data at four spatial resolutions (red: 1.25 mm iso., green: 1.5 mm iso., blue: 2 mm iso., yellow: 2.5 mm iso.) display the distribution (e.g., median, interquartile range, and range) of SW from 20 subjects in the upper panel. The lower table lists the relative difference of SW at lower resolution compared to the SW at higher resolution (each row) for different tractography methods (each column), with asterisks denoting significance levels (*: $p < 0.05$, **: $p < 0.01$, ***: $p < 0.001$). The color of each table cell indicates the magnitude and direction of the SW difference (blue: decrease, red: increase)
